# Protein context shapes the specificity of domain-peptide interactions *in vivo*

**DOI:** 10.1101/2020.05.18.103002

**Authors:** Ugo Dionne, Émilie Bourgault, Alexandre K Dubé, David Bradley, François JM Chartier, Rohan Dandage, Soham Dibyachintan, Philippe C Després, Gerald D Gish, Jean-Philippe Lambert, Nicolas Bisson, Christian R Landry

## Abstract

Protein-protein interactions (PPIs) between modular binding domains and their target peptide motifs are thought to largely depend on the intrinsic binding specificities of the domains. By combining deletion, mutation, swapping and shuffling of SRC Homology 3 (SH3) domains and measuring their impact on protein interactions, we find that most SH3s do not autonomously dictate PPI specificity *in vivo*. The identity of the host protein and the position of the SH3 domains within their host are both critical for PPI specificity, for cellular functions and for key biophysical processes such as phase separation. Our work demonstrates the importance of the interplay between a modular PPI domain such as SH3 and its host protein in establishing specificity to wire PPI networks.

## Introduction

Proteins often display a modular architecture defined by folded domains that bind short linear peptide motifs on their interaction partners (*1*). Modular domains are generally considered to act as “beads on a string” by virtue of their ability to independently fold and bind target peptides with high intrinsic specificity *in vitro* (*2*–*5*). However, binding domains are often part of larger proteins that can comprise many functional elements. Whether/how PPI domain binding specificity is modulated by positioning within their host protein and/or intramolecular interactions (collectively defined here as protein context) remains poorly defined. Such regulation of interaction specificity would imply that during evolution and in disease states, mutations occurring either within or outside a modular binding domain could alter its protein interaction specificity. We examined this question by studying SRC Homology 3 domains (SH3s) PPIs *in vivo*. SH3s are one of the most prevalent families of modular binding domains, having expanded in number throughout evolution with 27 in yeast (on 23 proteins) and nearly 300 in human (*6*–*8*). These ~60 amino acid domains are present on signaling proteins, regulating functions such as endocytosis and actin cytoskeleton remodelling (*9*). SH3s typically bind to Pro/Arg-rich peptide motifs on their target partners with an archetypical PXXP motif (where X represents any amino acid) (*7*, *10*). In order to fully understand how PPI networks achieve specificity and how it is altered by mutations, domain gains and losses, we combined genome editing, cellular phenotyping and proteomics to determine the *in vivo* contribution of protein context to SH3 domain specificity and functions.

### SH3s contribute to PPI networks complexity and protein function *in vivo*

To assess the requirement of SH3 domains for PPIs, we first measured binary interactions between 22 yeast WT SH3-containing proteins (Fig. S1A) as baits and 575 putative partners from their interconnected signaling networks using the dihydrofolate reductase protein-fragment complementation assay (DHFR-PCA) in *S. cerevisiae*. About 33% (202/607) of the detected PPIs were described before, mostly by direct methods (Fig. S1B-C) (*11*). We repeated the experiments with baits in which the SH3s were individually replaced with a flexible linker by genome editing (SH3 domain deletion/stuffing, Fig. 1A). About a third of the SH3-containing protein interactome is qualitatively or quantitatively SH3-dependent (171 PPIs out of 607, Fig. 1B and Fig. S1B-C). We validated the vast majority of the quantitative changes in low-throughput experiments and excluded that this is simply due to changes in protein abundance (Fig. S1D, E and F). SH3 binding motifs (*12*) are more frequent in the SH3-dependent PPI partners when compared to sequences from random proteins, SH3-independent PPI partners or PPI partners that are stronger or gained following SH3 deletions (p-values = 1.5×10^−17^, 0.0038 and 2.1×10^−08^, respectively; Mann-Whitney test, one-tailed, Fig. 1C). The changes we measure are therefore enriched for PPIs that depend on the direct interactions of the domains with the PPI partners. Interestingly, randomly assigning SH3 binding motifs does not significantly affect the motif enrichment observed (p=0.25, Mann-Whitney test, two-tailed), indicating that the SH3-specific peptide motifs determined *in vitro* (*12*) can partially identify SH3-bound proteins but poorly discriminate among different SH3 domains *in vivo* (Fig. 1D).

**Fig. 1.**
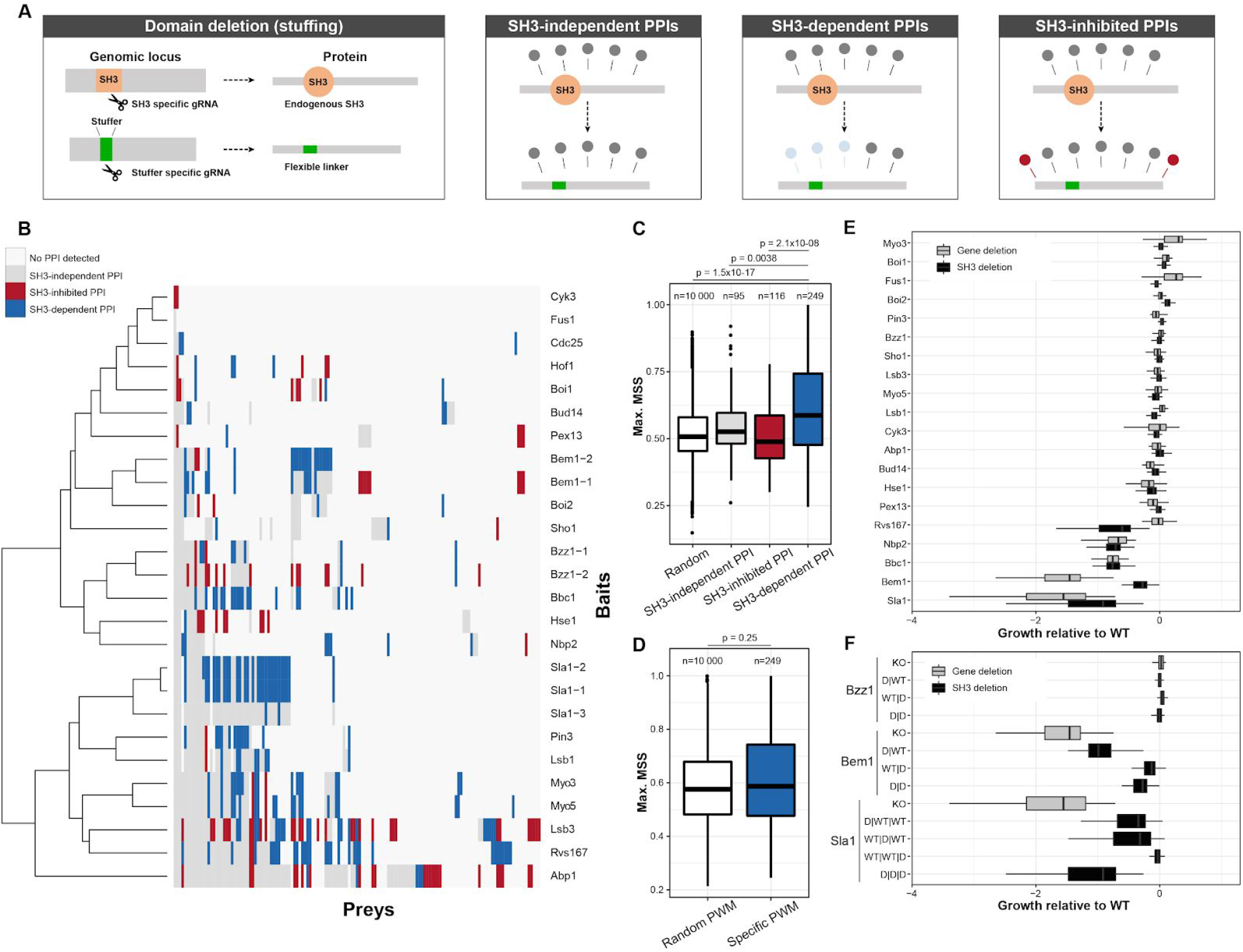
Definition of SH3-dependent PPIs *in vivo* and their functional impact. A) CRISPR-Cas9 SH3 editing approaches to study SH3 domains in living cells. For domain deletion (stuffing), SH3 sequences are replaced by a stuffer sequence encoding a linker using CRISPR-Cas9. SH3 dependency can then be tested. B) PPIs of WT and SH3-deleted proteins. Colors represent different types of PPIs (assessed in quadruplicate). C) Matrix similarity scores (MSS) of SH3-specific Position Weight Matrices (PWMs) for preys corresponding to different types of PPIs from B). SH3-dependent PPIs are enriched for SH3 binding motifs relative to the Random, SH3-independent, and SH3-inhibited PPIs (p = 1.5×10^−17^, p = 0.0038 and p = 2.1×10^−08^, Mann-Whitney test, one-tailed). D) PWM MSS for sequences of SH3-dependent preys (as in 1C), and for PWMs randomly assigned to SH3s (p = 0.25, Mann-Whitney test, two-tailed). E) Growth of gene- or SH3-deleted strains compared to WT under 51 stress conditions. For Bzz1, Bem1 and Sla1, the data shows deletion of all SH3s. F) Same as E) but for individual or combinations of SH3 deletions. D represents SH3 replaced with a linker sequence. Strains were grown in twelve replicates (E-F). Data is in Table S2, including the names of the preys in panel B, and in Table S1.

Surprisingly, 37 PPIs are increased and 75 are gained following SH3 deletions (Fig. 1B). Some of these PPIs have previously been detected with other methods (Fig. S1B-C). It is therefore likely that SH3 deletion alters protein folding or positioning within a complex or its relative binding preference, making these gained PPIs now detectable by DHFR-PCA. This observation may also be explained by changes in intramolecular interactions, as reported for the human SH3-containing SRC-family of tyrosine kinases (*13*).

We next assessed the contribution of SH3s to cellular phenotypes by measuring the growth of WT, SH3-deleted and knockout strains under 51 different stress conditions ranging from DNA damage induction to high osmolarity (Fig. 1E-F). In most cases, knockout or SH3 deletion leads to subtle phenotypes (Fig. 1E), as expected for non-essential genes (*14*). When gene deletion results in strong phenotypes, SH3 deletion consistently leads to similar growth defects (e.g. *NBP2, BBC1, BEM1* and *SLA1*). The number of SH3-dependent PPIs of a SH3-protein is negatively correlated with the growth score of the SH3-deleted strain relative to its WT strain (Pearson’s correlation = −0.426, p-value = 0.038). These results highlight the critical role of SH3 domains and their PPIs to protein function.

### SH3s are rarely sufficient to dictate the breadth of PPIs driven by their host

Having determined that multiple PPIs require SH3s, we asked whether the latter are autonomously capable of establishing PPI specificity in their host proteins by swapping domains (Fig. 2A). Using a second round of genome editing (*15*), we individually replaced Abp1 SH3 with the 27 yeast SH3s and 3 human Abp1 orthologs’ SH3s (CTTN, HCLS1 and DBNL, Fig. 2B). The reintroduction of its own SH3 (Abp1_SH3_ in Abp1) reconstitutes almost perfectly Abp1’s interaction profile (Kendall’s *τ* = 0.93, Fig. 2B). For cases where Abp1 loses most of its PPIs, Abp1 expression level was not significantly affected by SH3 swapping (Fig. S2A). No homologous SH3 domain re-establishes the normal Abp1 PPI profile (Fig. 2B and Fig. S2B). However, we observe a significant correlation (cophenetic correlation = 0.20, p-value = 0.005, Fig. 2C) between the similarity of PPI profiles (Fig. 2B) and the sequence similarity of the SH3s. For instance, Abp1 swapped with SH3s from its human orthologs, which have the highest sequence identity (human orthologs 46 to 52%, other yeast SH3s 19 to 41%), displays PPI profiles most strongly correlated with WT Abp1 (Kendall’s *τ:* CTTN = 0.89, DBNL = 0.89 and HCLS1 = 0.86, Fig. 2B and Fig. S2B). Nonetheless, a subset of SH3-dependent PPIs is only observed with the endogenous domain. For instance, Hua2 and App1, both well characterized partners of Abp1 (*12*, *16*, *17*), are only detected when Abp1 contains its own SH3 (WT or swapped via editing). A given SH3 is therefore not fully replaceable with other paralogous or orthologous domains. This pattern is confirmed by cellular growth phenotypic analyses (Fig. S2C-D). PPIs, SH3 sequence similarity and growth profiles under stress conditions are correlated (Fig. S2C-D). This relationship is particularly clear when analyzing growth in hygromycin. Indeed, *ABP1* deletion leads to hygromycin resistance (*18*) and this phenotype is dependent on Abp1 SH3 (Fig. S2D). None of the swapped SH3s, except for the Abp1_SH3_ in Abp1, fully reproduces WT sensitivity, confirming our observation with PPI patterns (Fig 2B and Fig. S2D). However, the orthologous human SH3s CTTN_SH3_, HCLS1_SH3_ and DBNL_SH3_ in Abp1 display intermediate phenotypes; this is consistent with PPI profiles and sequence similarity clusters (Fig. 2B-C and Fig. S2D).

**Fig. 2.**
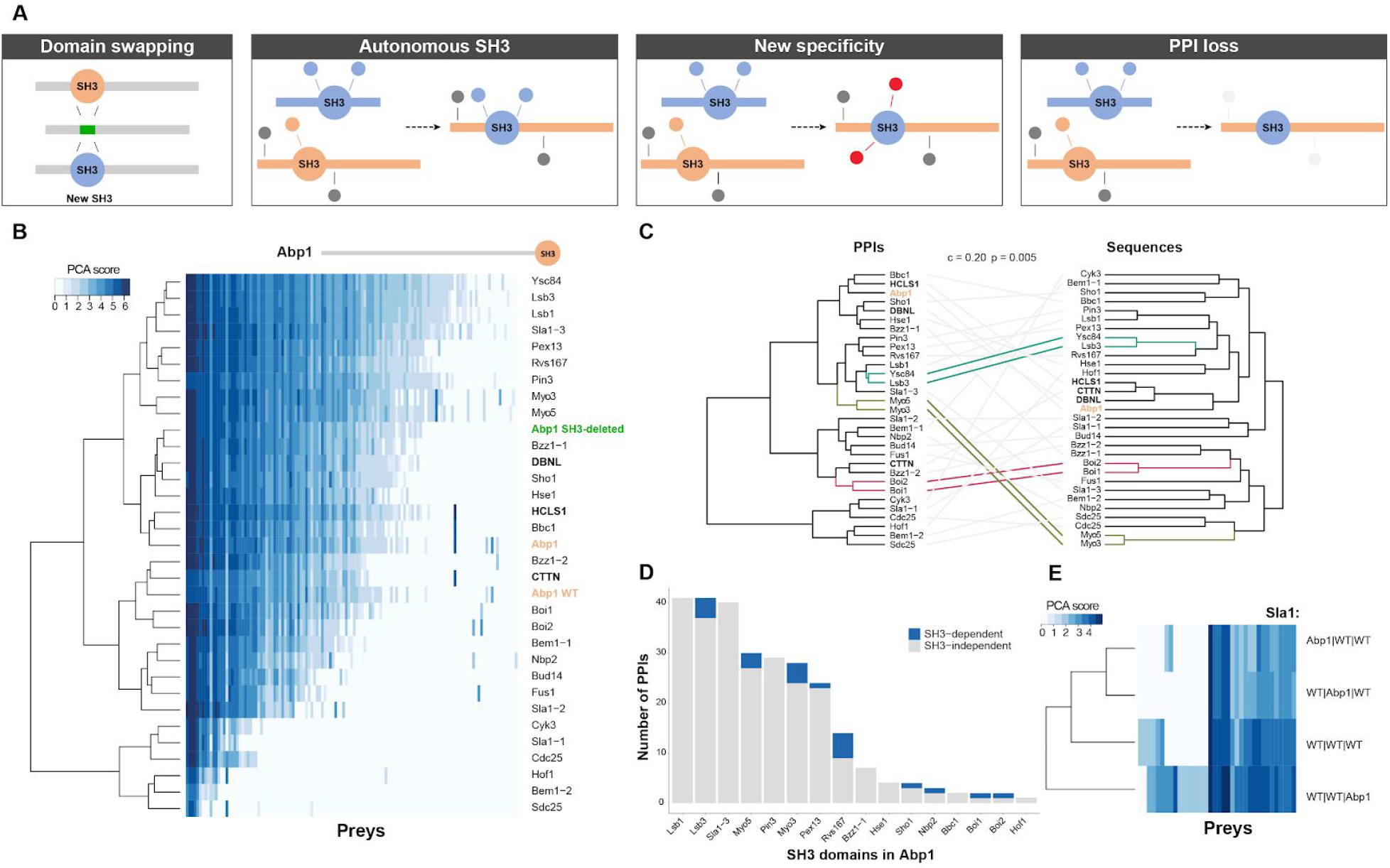
SH3 domains are rarely sufficient to establish their endogenous specificity in their host protein. A) *In vivo* SH3 domain swapping. The stuffer DNA from a SH3-deleted strain is replaced by SH3s from other genes. Consequences of SH3 swapping on the host protein PPIs are illustrated. B) PPIs of Abp1 with its endogenous SH3 domain or with a SH3 from other yeast proteins or from its human orthologs. Blue shades correspond to PPI interaction scores measured by DHFR-PCA (PCA score). A scaled cartoon representation of Abp1 is shown above the heatmap. Bold SH3 domains represent human orthologous, orange SH3s are controls, and green is for the SH3-deleted protein. Each PPI was assessed in quadruplicate. C) Cophenetic correlation between Abp1 SH3 swapped interaction clusters with the SH3 domain sequence similarity clusters. The empirical p-value (p = 0.005) was obtained from permutation. D) Number of PPIs gained by Abp1 upon domain swapping. PPIs originally detected as dependent on the domain when present in its host protein are shown in blue. E) Sla1 PPIs with the SH3 from Abp1 inserted at each of the three SH3 positions. Each SH3 is either the WT Sla1 domain (WT|WT|WT) or is Abp1 SH3 (Abp1|WT|WT is Sla1 with its SH3-1 swapped with Abp1 SH3). PPIs were measured in quadruplicate. See Table S3 for B-D), and Table S5 for E).

Several SH3 swappings lead to an inhibition of >50% of Abp1 PPIs, including SH3-independent interactions (Fig. 2B and Fig. S2B). This suggests that SH3s can affect binding that is mediated by other regions of the protein, most likely through allosteric effects. In addition, two thirds (21/30) of SH3 swaps lead to gains of PPIs that were not detected with WT Abp1 (Fig. 2B Fig. S2B). As expected, some SH3s can bring a subset of their SH3-dependent PPIs to Abp1 protein context; for example, Sho1_SH3_ and Nbp2_SH3_ in Abp1 promote the interaction with Pbs2 (*19*–*21*) (Fig. 2D Fig. S2E). However, the majority of the gained PPIs were not identified in our SH3 deletion screen as being SH3-dependent (Fig. 2D). Thus, most SH3s do not autonomously establish their endogenous specificity into a new protein context (Fig. S2E). Consistent with these results, partners gained by Abp1 SH3-swapped proteins are not enriched for SH3-specific binding motifs relative to unaffected Abp1 PPIs (p=0.52, Mann-Whitney test, one-sided, Fig. S2F).

We also examined whether the ability of a given SH3 to mediate PPIs depends on its position within the same host protein. We individually swapped Abp1 SH3 into each of Sla1’s three SH3 positions (Fig. 2E). Inserting Abp1 SH3 at either of the first two positions only slightly affects Sla1 PPIs (Fig. 2E). SH3 swapping at the third position led to the detection of seven new PPIs despite our observation that only two Sla1 partners depend on this third SH3 (Fig. 1B and Fig. 2E). None of the PPIs gained by Sla1 following Abp1 SH3 swapping were originally found to be dependent on Abp1 SH3, further supporting our finding that the ability of a domain to dictate its host PPI partners is highly dependent on the identity of the host and the position of the SH3. Overall, the observation that SH3s are rarely autonomous in dictating their host PPI partners, but rather alter PPIs in a manner that cannot be predicted from their intrinsic specificity suggests the presence of complex interactions between a SH3 domain and its host protein.

### Allosteric effects of SH3 domains on SH3-independent interactions

The above analysis revealed that domain swapping impacts PPIs in a sequence-dependent manner, with divergent SH3 domains having the strongest effects. Swapping SH3s into a protein also affects PPIs that were not previously found to be SH3-dependent, suggesting that SH3s can alter PPIs allosterically. To systematically investigate the distinction between effects on SH3-dependent and SH3-independent PPIs, we measured binding of Abp1 to a SH3-independent (Lsb3) and to a SH3-dependent partner (Hua2) for all possible single mutants of Abp1 SH3 (Fig. 3A-D and Fig. S3A-C). Mutation sensitivity profiles for the two targets are overall highly correlated (Kendall’s *τ* = 0.59, p-value = 1.0×10^−201^), but also show significant differences (Fig. 3B-D). These results are reproducible on a small scale (Fig. S3D), and confirm that SH3-independent PPIs (e.g. Abp1-Lsb3) are affected by changing the sequence of Abp1 SH3 (Fig. 3C).

**Fig. 3.**
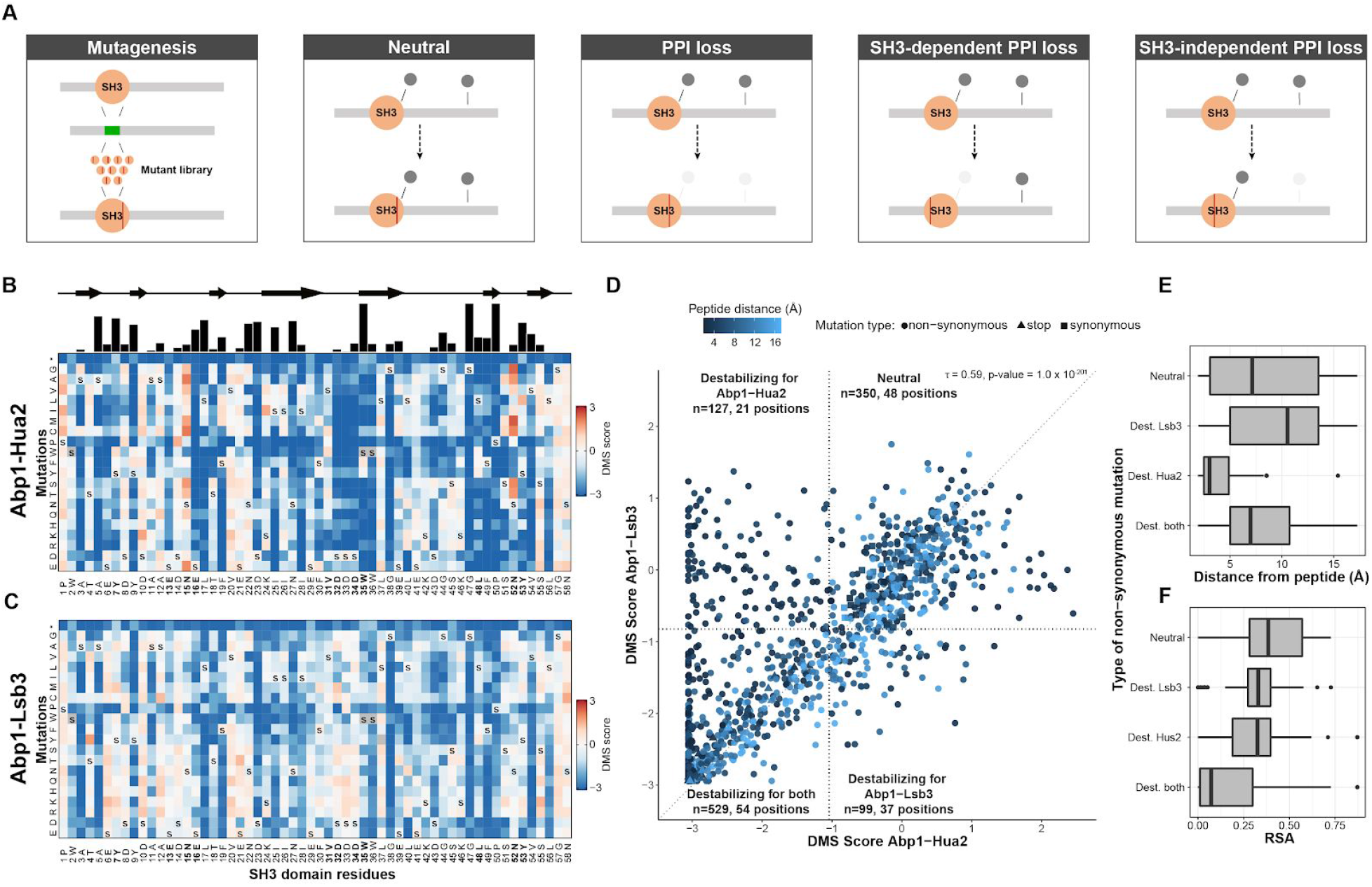
SH3 domain sequences affect both SH3-dependent and -independent PPIs. A) The stuffer DNA of a SH3-deleted strain is replaced by a library of single mutants. Categories of mutations based on their effect on PPI types are shown. B-C) Impact of mutations on binding represented in terms of Deep Mutational Scanning (DMS) score. B) Tolerance to SH3 mutations of the SH3-dependent interaction Abp1-Hua2. Blue: low DMS score i.e. reduced interaction strength, red: increased strength. Secondary structure of the SH3 is shown above. The black bars on the top represent the level of conservation across the 27 *S. cerevisiae* SH3s. Positions in bold represent residues in contact with the ligand (see also Fig. S3F). C) Same as B) but for the SH3-independent Abp1-Lsb3 PPI. The data shows the average of two biological replicates. D) DMS scores for the two PPIs. DMS scores below the 1st percentile of synonymous variant (dashed line) are defined as destabilizing the PPI. Four categories of mutants are shown: (1) destabilizing both PPIs, (2) specifically destabilizing Hua2 PPI or (3) Lsb3 PPI and (4) neutral or slightly increasing binding. The minimum distance relative to a binding peptide is shown in blue (Ark1 peptide, (*22*)). E) Minimum distance to the Abp1-bound peptide and (F) the relative solvent accessibility (RSA) for each category. See also Table S4. Dest is for destabilizing mutations.

Some positions are sensitive to any mutation for both PPI types (e.g. L17), while others are specific to one (e.g. D32). Mutations affecting both PPIs correspond to buried residues that are distal from the Abp1-bound peptide and likely affect protein folding rather than amino acid interactions (*22*) (Fig. 3D-F and Fig. S3E). Positions that are only destabilizing the Abp1-Hua2 lie at the peptide interface, which is in clear contrast with the sensitive positions that specifically destabilize Abp1-Lsb3 and that are distant from the SH3 binding peptide (Fig. 3D-F and Fig. S3F). Few positions specifically affecting the Abp1-Hua2 SH3-dependent interaction are conserved between Abp1 SH3 and other yeast SH3s; this could explain why none of the other SH3s can complement its loss upon domain swapping (Fig. 2B and Fig. 3B). A subset of mutations at positions 15 and 52, both predicted to be in contact with a target peptide, specifically strengthen the Abp1-Hua2 interaction, which could in principle have a higher affinity (Fig. 3B, C and Fig. S3F).

Overall, this analysis helps discriminate the residues defining SH3 binding specificity from the positions regulating the core functions of the domain. The latter most likely interact in an allosteric way with the binding of partners that do not depend on the intrinsic binding specificity of the domain.

### SH3 positions are not interchangeable in multi-SH3 proteins

Proteins containing multiple SH3s can mediate the formation of multivalent interactions, bringing an additional level of complexity to PPI regulation. The position of SH3s in their host is highly conserved and is generally independent of the extent of their amino acid sequence conservation (Pearson’s correlation = 0.22, p-value = 0.28, Fig. S4A-C), suggesting that SH3 positioning is key to function. To quantify the importance at the network level of SH3 position in their host, we focused on Sla1, a cytoskeleton binding protein that has three SH3s with low sequence identity (Fig. 4A, B and Fig. S4A).

**Fig. 4.**
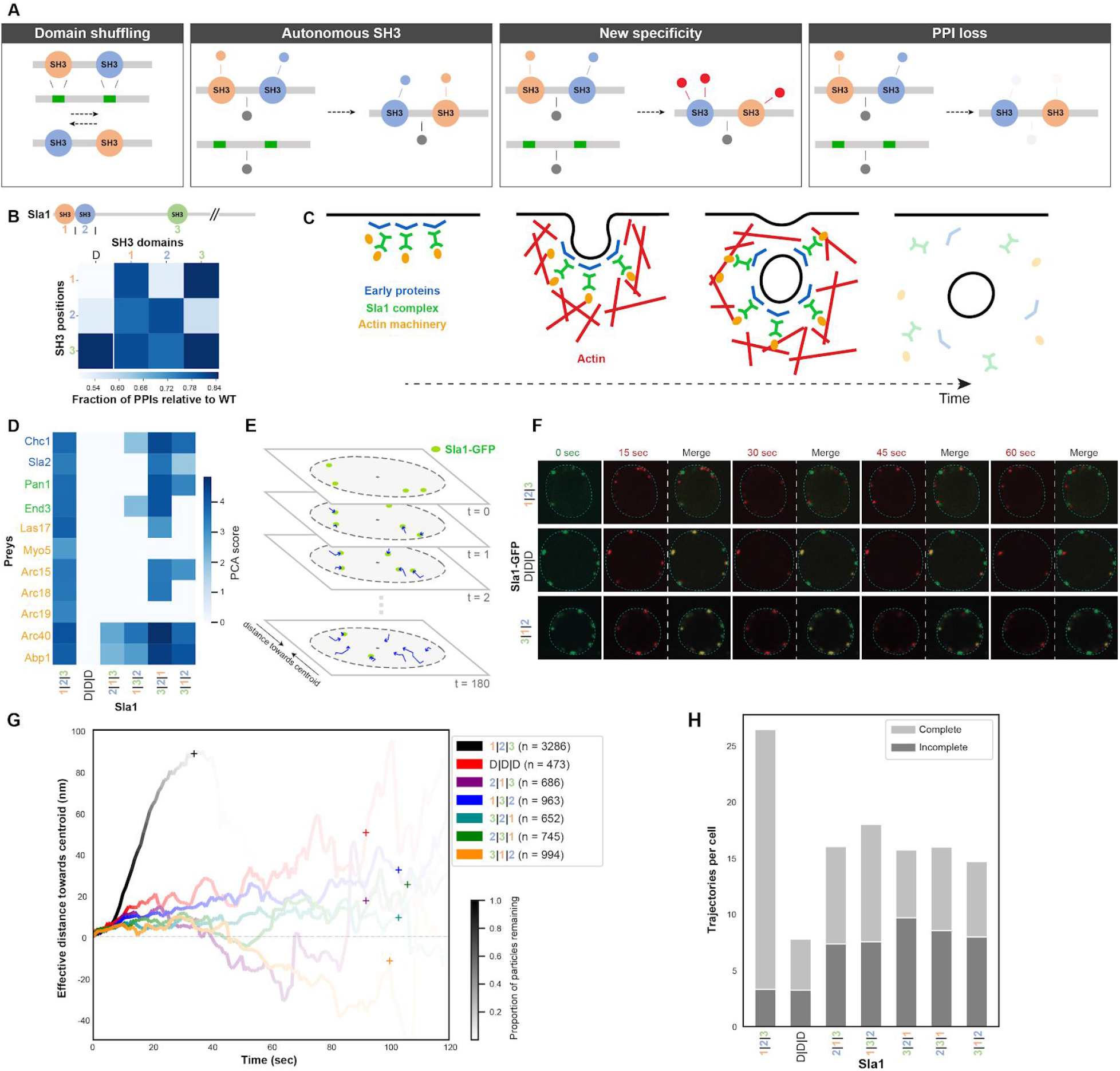
Shuffling SH3 positions alters Sla1 interactome *in vivo* and impacts its function in endocytosis. A) Domains are moved from one position to another in proteins containing multiple SH3s (domain shuffling). Possible outcomes on PPIs are presented. B) Number of PPIs detected relative to Sla1 WT for SH3-shuffling or -deletion per SH3 position (measured in quadruplicates). C) Function of Sla1 as an adaptor linking early proteins with the actin machinery in clathrin-mediated endocytosis. D) PPIs of Sla1 SH3-shuffled or -deleted with clathrin-mediated endocytosis related partners. The color code represents the strength of the PPIs (PCA score). Sla1 1|2|3 represents the WT protein. E) Schematic of Sla1 foci assembled at the cell membrane and their movement toward the center of the cell during internalization before disassembly, in time. F) Representative microscopy images of cells expressing different Sla1-GFP proteins at multiple time points. Foci from WT Sla1-GFP, the negative control D|D|D and a Sla1-GFP shuffle (3|1|2) are shown. Green: first time frame; red: others. A merge with the starting point for each time frame is shown. G) Average effective distance travelled by Sla1-GFP particles towards the centroid of the cell for the different constructs. Color transparency represents the proportion of events that are not completed yet. Plus signs (+) represent the moments in time when 95% of the foci have disassembled. H) Average number of complete or incomplete events, i.e. that were not completed before the end of the image acquisition period. See also Table S5.

Significant differences in PPIs and growth phenotypes are dependent on Sla1 SH3 domains (Fig. 1B and 1F). We constructed all possible domain-position permutations within Sla1 (i.e. domain shuffling, Fig. 4A). Sla1 PPIs are highly dependent on SH3 positions (Fig. 4B and Fig. S5A). As Sla1 SH3-1 and SH3-2 bind to the same peptide motifs *in vitro* (*12*), we expected little impact from exchanging their position if peptide recognition was the sole determinant of SH3 specificity *in vivo*. Surprisingly, shuffling the first two domains (2|1|3) results in the loss of ~80% of Sla1 PPIs (Fig. S5A). Many PPIs are lost when the SH3-1 position is occupied by SH3-2 (2|2|3) but maintained with SH3-3 (3|2|3), even if there are few PPIs that depend on SH3-3 when its two first endogenous SH3 are present (Fig. 1B and Fig. 4B). Shuffling two or all three Sla1 SH3s leads to the loss of most of Sla1 PPIs, despite all three SH3s still being present albeit in a different order (Fig. S5A). In general, the similarity of PPI profiles of the mutants is significantly correlated with their growth phenotypes (cophenetic correlation = 0.28, p-value = 0.00002, from permutation, Fig. S5B).

During clathrin-mediated endocytosis, Sla1 is an adaptor linking cargos to clathrin and recruits members of the actin machinery via SH3-dependent PPIs (Fig. 4C) (*23*, *24*). Sla1 compartmentalizes into foci at internalization sites on the plasma membrane where it is present until vesicles are fully internalized (Fig. 4C); its deletion was shown to alter endocytosis dynamics (*23*–*25*). The shuffling of Sla1 SH3 alters PPIs with partners involved in different phases of endocytosis (Fig. 4D). We followed mutant Sla1 alleles by measuring the distance traveled by Sla1-GFP-labeled vesicles from the cell periphery towards the cell center (Fig. 4E and Fig. S5C). Sla1-GFP foci movement is mostly regulated by Sla1 two N-terminal SH3s, as previously reported (Fig. S5D) (*24*, *26*). Shuffling Sla1 SH3s in any combination results in a drastic decrease in the effective distance traveled by Sla1-positive foci, as well as their persistence in time, likely due to the inability to complete the process (Fig. 4F-G and Fig. S5E-G). This observation is consistent with a >50% decrease in the number of Sla1-GFP-labeled particle internalizations per cell and an increase in the number of incomplete events (Fig. 4H and Fig. S5E). The linearity of Sla1-GFP foci trajectories is also drastically perturbed in all Sla1 SH3-shuffled strains, suggesting inefficient vesicle internalization (Fig. S5F). The identity and position of each of Sla1 SH3 domains therefore regulate its PPIs and quantitatively impact the cell’s ability to tolerate stress, and to perform dynamic processes such as endocytosis.

### Protein context influences human SH3s PPIs

We extended to human the study of multi-SH3 proteins, as they are more prevalent (62/216 SH3-containing proteins) than in yeast (3/23). NCK adaptors (NCK1 and NCK2) connect growth factor receptors to the actin machinery via their single SH2 and three SH3 domains (*27*), which bind to similar types of peptide motifs but are known to interact with different partners (*7*, *28*) (Fig. 5A). We combined affinity purification with sequential window acquisition of all theoretical mass spectra (AP-SWATH) quantitative proteomics to test whether SH3 shuffling impacts NCK2 PPIs in human cells (*29*, *30*).

**Fig. 5.**
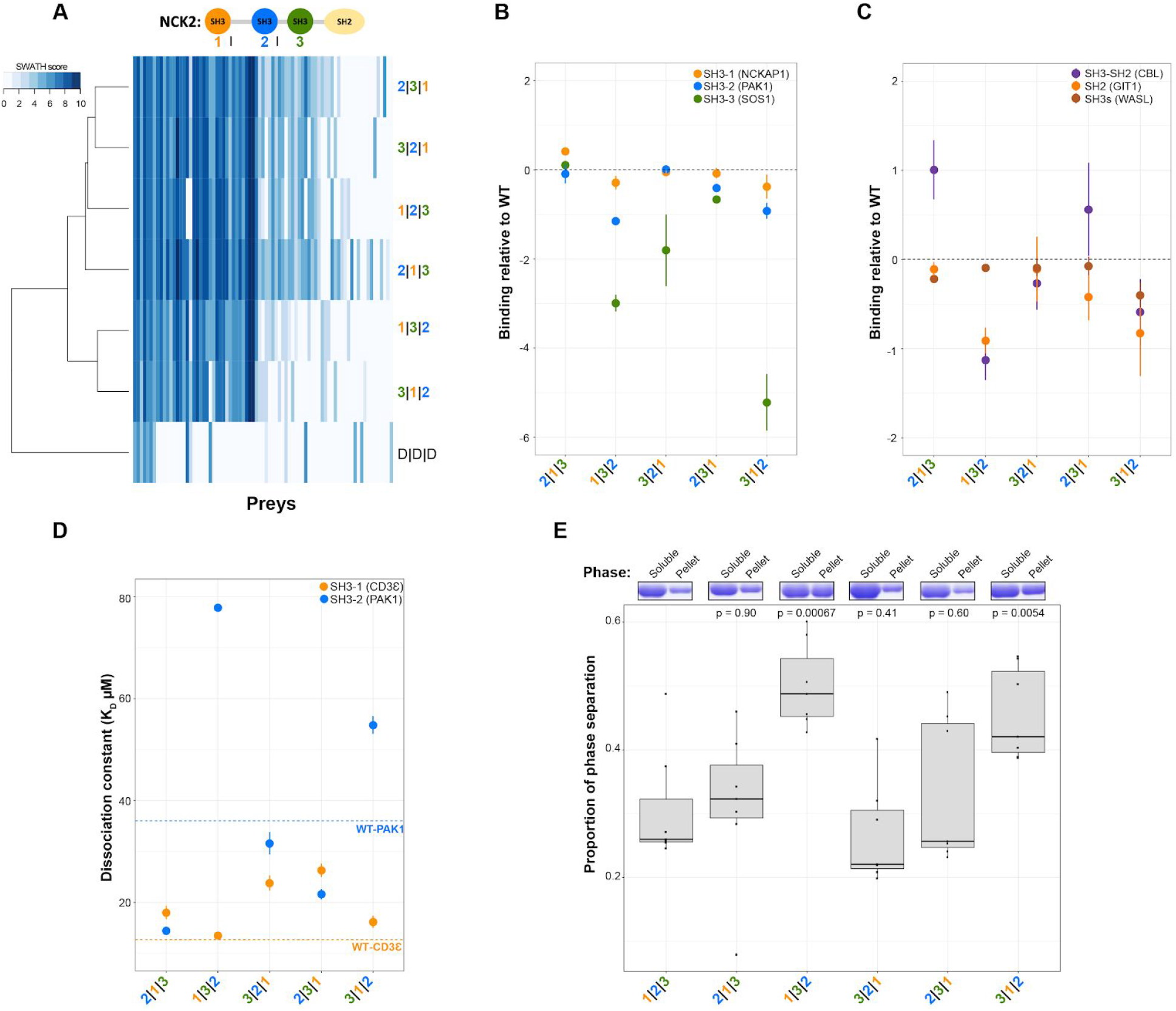
Human NCK2 SH3 domain shuffling alters its interactome in cells and its ability to phase separate. A-C) AP-SWATH quantitative MS data of NCK2 PPIs (duplicates, PPIs with a SAINT analysis FDR<1%, see methods). A) Scaled cartoon representation of NCK2 with its domains. Log2 of the average spectral counts (SWATH score) for the PPIs are shown. Baits were clustered based on the similarity of their PPI profiles. The color code represents the PPI strength (SWATH score). The bait nomenclature 1|2|3 is for WT NCK2 and D|D|D is for the triple SH3-inactive negative control (W38K/W148K/W234K). B-C) NCK2 PPIs for partners with known binding sites on NCK2. The spectral count from the two biological replicates was compared to the WT NCK2 score (Ratio NCK2mut/NCK2WT). The log2 average ratio is shown (WT NCK2 ratio = 0). Error bars indicate the SD. D) Fluorescence polarization dissociation constants (K_D_) for NCK2 full length recombinant proteins with a SH3-1 direct (CD3Ɛ) or SH3-2 direct (PAK1) partner (triplicates, K_D_ error values represent the SE). E) NCK2 phase separation after 24 hours of incubation. Soluble and phase-separated (pellet) proteins were quantified via Coomassie staining (typical replicate shown above). Proportion of proteins in the pellet compared to the total protein content (soluble + pellet) is shown for the seven replicates. Proportions of phase separated proteins were compared to NCK2 WT (pairwise ANOVA, p-values: 2|1|3 = 0.90, 1|3|2 = 0.00067, 3|2|1 = 0.41, 2|3|1 = 0.60 and 3|1|2 = 0.0054). See also Table S6.

We generated a high confidence WT NCK2 interactome (56 PPIs, FDR < 1%) that significantly overlaps with previously reported PPIs (71%), displaying about a third of all reported NCK2 interactors (*11*). We detect significant changes in the SH3-shuffled NCK2 complexes including the complete loss of up to 17 PPIs (30% of WT NCK2 PPIs) and the gain of up to 13 new partners (Fig. 5A). The SH3-3 position appears to be more critical as NCK2 PPI profiles cluster based on the identity of the SH3 at this location (Fig. 5A). Well-characterized SH3-dependent PPIs of NCK2 depend on the position of each target cognate SH3 (Fig. 5B). In particular, NCK2 SH3-2 association with PAK1 (*31*) is significantly impaired when the domain is at the third position (Fig. 5B and Fig. S6A-B). In contrast, NCK2 interaction with WASL, which is mediated by multiple SH3-PXXP interactions likely involving all three NCK2 SH3s (*32*), is only slightly altered by shuffling (Fig. 5C). SH3 shuffling also leads to longer-range disruptions of NCK2 SH2-dependent PPIs. For example, GIT1 and P130CAS/BCAR1 associations with NCK2 SH2 are significantly modulated in several SH3-shuffled mutants (*33*, *34*) (Fig. 5C and Fig. S6C-D). These results indicate that SH3 positioning can affect PPIs that are mediated by modular domains other than SH3s. Such indirect regulation of non-SH3 domains by adjacent SH3 positioning may also explain why many SH3-independent PPIs were altered by domain shuffling and swapping in yeast experiments.

To test whether shuffling affects the intrinsic SH3 affinity to targets via a direct effect on peptide binding, we performed *in vitro* fluorescence polarization binding assays using purified full length recombinant NCK2 and peptides from direct SH3 partners CD3Ɛ (SH3-1) (*35*) and PAK1 (SH3-2). SH3-1 shuffling leads to modest variations for NCK2 interaction with CD3Ɛ, with a maximum of ~2-fold decrease when NCK2 SH3-1 is moved to the third position (Fig. 5D and Fig. S6E). Similarly, NCK2 binding to its PAK1 target peptide decreases by ~2-fold when its SH3-2 is inserted at the SH3-3 site (Fig. 5D and Fig. S6F). These results indicate that shuffling SH3s in their host protein might only slightly affect the position and/or accessibility of their binding pocket even though changing the positions of NCK2 SH3s resulted in the complete loss of a subset of PPIs in cells (Fig. 5A). The consequences of SH3 shuffling on the NCK2 interactome are thus possibly enhanced by factors other than binding pocket availability *in vivo*.

### SH3 position regulate phase separation

The separation of phases in the cytosol is an important feature of multiple cellular processes. The ability of NCK2’s close paralog NCK1 to phase separate depends on both its number of functional domains and its capacity to self-associate (*36*). We examined the ability of NCK2 SH3-shuffled proteins to undergo phase separation *in vitro* via self-association. Remarkably, the different NCK2 SH3-shuffled mutants initiate varying levels of phase separation (Fig. 5E). The most striking differences occur when the SH3-2 is shuffled at the third position, which results in a significant ~2-fold increase in phase separation relative to WT (Fig. 5E). Phase separation is inhibited with the addition of a PAK1 peptide, presumably by competing homotypic binding to NCK2 SH3-2 (Fig. S7A). Based on this and previous observations on NCK1, we hypothesized that phase separation is dependent on NCK2 SH3-2 electrostatic interaction in *trans* with the first interdomain region of another NCK2 molecule (*36*), which would depend on salt concentration. The propensity of NCK2 SH3-shuffled proteins to phase-separate is altered by increasing NaCl concentration; constructs bearing the SH3-2 at the third position are the most resilient (Fig. S7B). In contrast, the addition of 1,6-hexanediol, a small aliphatic alcohol thought to impact the weak hydrophobic interactions (*37*), disrupts all NCK2 SH3-shuffled proteins in a similar manner (Fig. S7C). Taken together, these results demonstrate that NCK2 phase separation is highly dependent on the position of its SH3 domains. Shuffling NCK2 SH3-2 at the third position severely disrupts NCK2 PPIs in cells (Fig. 5A) while it stabilizes NCK2 condensates *in vitro* (Fig. 5E), suggesting that NCK2’s capacity to phase separate might impact its potential to nucleate PPIs *in vivo*.

## Conclusion

PPI networks dictate how signaling pathways and biological processes are physically organized (*38*). In human cells, it has been estimated that around half a million binary PPIs occur (*39*), and that a given individual might express nearly one million distinct proteoforms (*40*). It is therefore an outstanding challenge to understand how proteins find their cognate PPI partners while avoiding spurious interactions. Here, we demonstrate that most SH3s mediate specific interactions in combination with their host proteins. Our results suggest that the description of protein domains as “beads on a string” is too simplistic and that the protein context of structured domains is also of great importance. We assert that our findings may be extended to other families of protein binding modules, and therefore contribute to a better understanding of complex signaling networks in normal and disease states. Our observations also suggest that the evolution of protein domains by gains and losses can have complex effects that will depend on the sequential ordering of the domains themselves, and this beyond motif recognition residues and individual domain-peptide interactions.

## Acknowledgments

We thank Bisson and Landry lab members for discussions; Véronique L’Italien for expert assistance with figure artwork; N. Aubin-Horth, S. Elowe, S. Michnick, I. Gagnon-Arsenault and C. Gagné-Thivierge for critical reading of the manuscript. This work was supported by a Canadian Institutes of Health Research (CIHR) Foundation grant 387697 and a HFSP grant (RGP0034/2018) to CRL, CIHR Project grant 162439 to NB, and Discovery Grant 1304616-2017 from the Natural Sciences and Engineering Research Council of Canada (NSERC) to JPL. CRL holds the Canada Research Chair in Evolutionary Cell and Systems Biology. NB holds the Canada Research Chair in Cancer Proteomics. JPL is supported by a Junior 1 salary award from the Fonds de Recherche du Québec-Santé (FRQ-S). U.D. holds an Alexander-Graham-Bell Canada graduate scholarship and the Louis-Poirier scholarship of the Fondation du CHU de Québec-Université Laval. EB was supported by a NSERC graduate scholarship. RD is funded by a FRQ-S Postdoctoral fellowship. SD was supported by a MITACS globalink scholarship. PD is supported by a CIHR Vanier Canada Graduate Scholarship.

## Author Contributions

UD, AKD, FC, RD, NB and CRL designed experiments. UD, EB, AKD, RD, SD, FC and GDG performed experiments. DB, PD, RD, SD and CRL designed and performed computational analyses. UD, EB, DB, RD, PCD, CRL, and JPL analysed results. UD, NB and CRL wrote the manuscript with contributions from all authors. NB and CRL supervised the research.

## Materials and Methods

### *Saccharomyces cerevisiae* strains

For the DHFR-PCA experiments, bait strains were constructed and preys were either constructed or retrieved from the yeast protein interactome collection (Horizon) (*42*). BY4741 (MATα, *his3Δ leu2Δ met15Δ ura3Δ*) strains with a specific gene of interest fused at its 3’ to the DHFR F[1,2] with a nourseothricin-resistance marker (NAT 100 μg/ml, Werner BioAgents) were used as baits. BY4742 (MATα, *his3Δ leu2Δ lys2Δ ura3Δ*) prey strains are each expressing a gene of interest fused at its 3’ end to the DHFR F[3] and are resistant to hygromycin B (HPH 250 μg/ml, Bioshop Canada). The same BY4741 strains were used in the growth assays with the addition of knockout (KO) strains from the yeast knockout collection (Horizon) (*14*). For the fluorescence microscopy experiments, the different Sla1 engineered proteins were constructed using the BY4741 Sla1-GFP strain from the GFP collection (Thermo Fisher Scientific) (*43*, *44*). See Table S7 for the complete list of all strains used for this study.

### *Saccharomyces cerevisiae* growth conditions

Cells were grown with their specific selection antibiotics in YPD medium (1% yeast extract, 2% tryptone, 2% glucose, and 2% agar (for solid medium)) and with the combination of NAT and HPH for diploid selection. Synthetic medium (0.67% yeast nitrogen base without amino acids and without ammonium sulfate, 2% glucose, 2.5% noble agar, drop-out without adenine, methionine and lysine, and 200 μg/ml methotrexate (MTX, Bioshop Canada) diluted in dimethyl sulfoxide (DMSO, Bioshop Canada)) was used for the DHFR-PCA experiments (PCA selection). For western blot experiments, cells were grown in synthetic complete medium (0.174% yeast nitrogen base without amino acids, without ammonium sulfate, 2% glucose, 0.134% amino acid dropout complete, 0.5% ammonium sulfate). Sla1-GFP strains were grown in synthetic complete medium without tryptophan to reduce background fluorescence. Cells were grown in synthetic complete medium with monosodium glutamate (0.1% MSG) instead of ammonium sulfate in hygromycin B growth experiments. See Table S7 for all the growth media used.

### Cells

Human embryonic kidney 293T cells (HEK293T, ATCC:CRL-3216) were grown in Dulbecco’s Modified Eagle’s medium (DMEM) high glucose complemented with 10% fetal calf serum under 5% CO_2_ at 37°C. Transient expression of cDNA in cells was performed by polyethylenimine (PEI, Millipore Sigma) transfections. Enrichment of phospho-dependent PPIs was done by treating HEK293T cells with the tyrosine phosphatase inhibitor pervanadate, freshly made by mixing 4 mM orthovanadate (Millipore Sigma) with 30% H_2_O_2_ (Millipore Sigma) in water in a 200:1 volume ratio.

### Strain construction

The complete list of strains, primers and gRNAs used in this study can be found in Table S7. All constructed yeast strains were validated by PCR and confirmed by Sanger sequencing of the locus of interest. Bait strains for the DHFR-PCA experiments were constructed as follows. The genes of interest were fused at their genomic 3’ end by replacing the stop codon with a linker (GGGGSGGGGS), the DHFR F[1,2] followed by a smaller linker (GGGGS), a 1X FLAG tag (DYKDDDDK) and the NATMX4 resistance cassette via homologous recombination (all amplified from pAG25-DHFR[1,2]-linker-FLAG). The set of 615 prey strains (DHFR F[3] fusions in the BY4742 background) were retrieved from the Yeast Protein Interactome Collection (*42*) with the exception of 73 that were reconstructed (Table S7). Knockout strains were taken from the yeast knockout collection except for *ABP1, BEM1, BZZ1* and *SLA1* which were reconstructed by replacing the ORFs with the NATMX4 resistance cassette by homologous recombination in the BY4741 background.

Yeast SH3 domain genomic sequences were replaced by CRISPR-Cas9 genome editing. Each SH3 domain (as defined by the SMART database V8.0 except for Ysc84 SH3 (as defined by PROSITE), see Table S1) was specifically targeted by one gRNA as described in (*45*). Yeast competent cells were co-transformed with a pCAS plasmid (Addgene plasmid 60847, (*46*)) expressing both the gRNA of interest and *Streptococcus pyogenes* Cas9 and a donor DNA sequence (stuffer) with 60bp homology arms surrounding the SH3 DNA sequence. The Stuffer DNA was either PCR amplified or digested from a plasmid. The stuffer DNA sequence (GGCGGAAGTTCTGGAGGTGGTGGT) codes for a small flexible linker (GGSSGGGG) inspired from protein linkers used in structural biology (*47*, *48*). This type of linker is frequently used for fusing proteins without affecting the structure of the linked proteins.

The stuffing of *YSC84* SH3 rendered the strain sterile and was therefore not used for this study as the assays require crosses. For genes containing two SH3s (*BZZ1* and *BEM1*), a combination of this first stuffer with a second one (GGTGGCTCAGGAGGAGGTGGTGGA) coding for a highly similar linker (GGSGGGGG) was used, which allowed targeting each SH3 position independently. The two N-terminal domains of *SLA1*, which are separated by six nucleotides, were stuffed by a single stuffer. Cells containing pCAS were selected via the G418 resistance cassette from the plasmid on YPD plates (YPD + NAT 100 μg/ml (DHFR F[1,2] selection) + G418 200 μg/mL (Bioshop Canada)). Colonies were randomly selected and grown in YPD media without G418 to allow plasmid loss. The correct insertion of the stuffer was validated by PCR and Sanger sequencing. The SH3-swapped, -shuffled and -mutated strains were constructed following a second CRISPR-Cas9 editing step with the same procedures. The stuffer sequences were targeted by specific gRNAs (one gRNA per stuffer) and replaced by PCR amplified specific SH3 (with 40bp homology arms) or libraries of mutated SH3 domains for Deep Mutational Scanning (DMS) with 78bp flanking homology regions. Validation of the SH3 domain insertions was done as we previously described except for DMS libraries (see below).

### Cloning

The fragment of plasmid pAG25-DHFR[1,2]-linker-FLAG was synthesized by Synbio Technologies. A GGGGS sequence was added after the DHFR F[1,2] followed by one repeat of the FLAG epitope (DYKDDDDK) in the parental plasmid pAG25-DHFR[1,2] (*42*). Cloning of the gRNAs in pCAS was performed as described in (*45*). The cloning of the gRNAs sequences was performed via full plasmid PCR amplification with the KAPA (Roche) polymerase using 60-mer oligonucleotides (Eurofins Genomics). The resulting PCR products were digested for 1h at 37°C with DpnI (NEB) and transformed in competent *Escherichia coli* MC1061 cells. Positive colonies were selected on 2YT plates (1% yeast extract, 1.6% tryptone, 0.2% glucose, 0.5% NaCl, and 2% agar) with 100 μg/ml of kanamycin. The SH3 domain DNA sequence of *ABP1* used in the DMS experiments was cloned into pUC19 (NEB) via restriction enzyme cloning with 78bp homology arms surrounding the SH3 DNA (from genomic DNA). The sequences of CTTN, DBNL and HCLS1 SH3 domains (gBlocks, Integrated DNA Technologies) were cloned into pUC19 in the same way. Positive clones were selected on 2YT plates with 100 μg/ml of ampicillin. For NCK2 constructs, the mouse mNCK2 cDNA sequence was cloned via restriction enzymes in frame with a N-terminal 3XFLAG tag in pcDNA3.1(-) (ThermoFisher Scientific) for western blot (WB) experiments and into the pMSCVpuro vector (Clontech) for mass spectrometry (MS) experiments. For bacterial recombinant protein expression and purification, the cDNA was inserted in frame with a 6xHis-GST TEV-cleavable tag into a modified version of pET-30b (Novagen). The NCK2 SH3 shuffled chimeras were constructed using NEBuilder HiFi DNA Assembly Cloning Kit (New England Biolabs, Inc). The different functional regions of NCK2 are based on NCBI (NP_035009.3) and Uniprot (O55033) definitions and defined as SH3-1 (2-59), linker 1 (60-113), SH3-2 (114-170), linker 2 (171-197), SH3-3 (198-256), Linker 3 (257-282), SH2 (283-380). All cloning was validated by Sanger sequencing.

### PCA experiments and analysis

The overall DHFR-PCA pipeline is based on (*42*) and the procedures have been previously described (*49*). The screens were performed with robotically manipulated pin tools (BM5-SC1, S&P Robotics Inc). First, the preys were cherry-picked from 96-well plates to omnitrays in a 384 format on YPD + HPH solid media. Plates were then combined into the 1,536 format on YPD + HPH. The array was constituted of 575 preys previously shown to interact physically with at least one SH3-containing protein as reported in BioGRID V3.4.16 along with 40 negative controls (*11*). The prey array comprised a border of control strains of two rows and two columns corresponding to the PPI between LSM8-DHFR F[1,2] and CDC39-DHFR F[3]. This double border was used to remove border effects. Preys are present in duplicate at a randomized position in the 1,536 array (except for YAL041W which is present four times per array). Baits were crossed with two prey plates, resulting in four independent bait-prey tests for each PPI. Baits were grown in liquid YPD + NAT and then transferred on solid YPD + NAT omnitrays. They were then replicated on YPD + NAT omnitrays in 1,536 array format. Mating of DHFR F[1,2] baits with DHFR F[3] preys was done on YPD followed by incubation at 30°C for 48 hours. Two diploid selection steps were performed by replicating the plates on YPD + NAT + HPH and incubating them for 48 hours at 30°C. Pictures were taken at the end of the second diploid selection for quality control. All images were acquired with a EOS Rebel T5i camera (Canon). Finally, the diploid cells were replicated on omnitrays containing solid PCA selection media and incubated for 4 days as a first selection step in a spImager custom robotic platform (S&P Robotics Inc). Cells were replicated for a second PCA selection step and incubated in the same manner. The last time point of the second PCA selection step was used for the data analysis as in previous studies using this method.

Colony sizes from pictures were measured with the R package ‘Gitter’ (*50*). The output was validated by manual inspection. Colonies that did not grow in the diploid selection were removed from the analysis. Diploids that were still present with at least two replicates were kept for downstream analysis. The colony sizes were log2-transformed and each plate was normalized to its background level to allow for comparison among plates. The median log2 normalized colony size was then calculated for each PPI (PCA score). A cut-off was determined based on the PCA score distribution of all PPIs and on the comparison with controls to identify true PPIs (see below). To be called as a true PPI, a pair of bait-prey needed to have the median of its replicates (PCA score) above the threshold (see below). None of the baits used in the DHFR-PCA screens were found to interact with the 40 negative control preys. The first DHFR-PCA screen included WT, SH3-deleted and Abp1 SH3 swapped strains. The plates Sla1_SH3-1_ in Abp1, in which Sla1 SH3-1 was used to replace Abp1 SH3, had no detectable level of background and was assigned the measured background of Abp1 WT plates. The threshold of this screen was determined based on the distribution of the PCA score of every PPI and corresponds to the top 7.3% of all detected colonies in the DHFR-PCA experiment. PPIs above the threshold of the Abp1 WT bait are highly similar (Kendall’s r = 0.93) to those of the Abp1_SH3_ in Abp1 control bait, which suggest that the determined threshold allows the detection of true interactions. In addition, we found that the true PPIs of the WT SH3-containing baits are enriched in PPIs that were previously reported (33.2%) when compared to the PPIs that did not pass the threshold (7.7% previously reported PPIs) (*11*). PPI detection was also highly reproducible between replicates (Pearson’s correlation = 0.91-0.93). The vast majority of the true PPIs of this screen are from four replicates (93.6%). To determine which PPIs were weaker or stronger following SH3 domain deletions, a ratio was calculated to compare the PCA scores of the PPIs of SH3-deleted baits with the PCA score of the WT baits (PCA score ratio). The same threshold was applied to the PCA score ratio, with the top 7.3% identified as stronger PPIs and the bottom 7.3% as weaker PPIs following SH3 deletions. The same principle was applied to identify PPIs that were stronger or weaker following domain swapping in Abp1. The PCA scores of the PPIs of the Abp1 SH3-swapped baits were compared to the PPIs of the Abp1_SH3_ in Abp1 bait.

Liquid DHFR-PCA low-throughput experiments were performed to validate the PPIs identified as affected by the SH3 domain deletions in the high-throughput solid DHFR-PCA screen (256 PPIs weaker or lost and 116 PPIs stronger or gained following domain deletions). Mating and diploid selection was performed as described for the large scale experiment, with the exception that the plates were in 384 format. Following the second diploid selection, diploid cells were inoculated in a 96 v-shaped well plate using a robotically manipulated 96 pin tool in synthetic complete medium with MSG at pH 6.0 with NAT and HPH. After 48h of growth, optical density (OD) was read using a TECAN Infinite F200 Pro and dilutions to 1 OD in sterile nanopure water were prepared. Cells were then diluted to 0.1 OD by combining 25 μl of cell suspension at OD = 1 and 225 μl of PCA selection media without agar in a 96 wells plate. Plates were incubated at 30°C in a TECAN Infinite M Nano and OD was monitored each 15 mins for 72 hours.

A separate DHFR-PCA screen was performed for the Sla1 SH3 shuffling experiment, with the same analysis as previously described. Three plates had no detectable level of background, which correspond to the second plate of the three Sla1 SH3 WT re-insertion controls. They were attributed the average of the background of the three first plates of the same baits. The threshold was also determined based on the distribution of the PCA scores of all PPIs and corresponds to the top 2.75% of all colonies detected in this experiment. The PPIs above the threshold of the Sla1 WT bait are highly similar (Kendall’s r = 0.89, 0.90 and 0.87) to those of the WT SH3 re-insertion control baits, indicating that the threshold allows the detection of true interactions. Almost all true PPIs are from four replicates (96.2%). The Sla1 SH3 shuffled strain SH3-2|SH3-3|SH3-1 DHFR F[1,2] was not successfully constructed and was excluded from the experiment.

### Growth assays in stress conditions

All WT, SH3-deleted, KO (except for *CDC25* which is the only essential SH3-containing gene in yeast), *ABP1* SH3-swapped and *SLA1* SH3-shuffled strains were grown on omnitrays in 1,536 format and robotically manipulated with a pin tool (the DHFR F[1,2] bait strains were used except for the KO strains). Each strain was randomly positioned in 6 replicates per plate and two plates per condition were performed, for 12 replicates per strain in total. A border of two rows and two columns of WT strains (BY4741) around the plate was used to remove border effects. Strains were grown on YPD media for two days at 30°C. They were then replicated on 51 different media that are described in Table S7 and were incubated at 37°C in a spImager custom platform with images acquired every two hours for four days. The colony sizes from plates pictures were also measured with the R package ‘Gitter’ and the output was manually inspected. Positions with abnormality in colony circularity were removed. The colony sizes after 74 hours of incubation at 37°C were log2 transformed and used for the analyses. First, the differences between the two plate replicates were corrected to remove plate effects. The median colony size was calculated for each strain per plate. Strains with a difference in their median colony size of more than 2 between the two plate replicates were removed. Finally, the average of the median colony size was calculated for each strain per condition (growth score).

For the hygromycin B resistance assays in liquid cultures, the *ABP1* SH3 swapped strains including the *ABP1* SH3-deleted and *ABP1* KO strain were grown overnight at 30°C as pre-cultures in synthetic complete medium with MSG. The next day, OD was measured with a TECAN Infinite F200 Pro plate reader and the cultures were diluted to OD of 1.0 in sterile nanopure water. The different strains were further diluted at 0.1 OD by adding 25 μl of cultures to 225 μl of synthetic complete medium with MSG or with MSG and Hygromycin B 120 μg/mL in 96 wells plate (final concentration of 108 μg/mL). Cells were incubated in a TECAN Infinite M Nano plate reader at 37°C for 24 hours and OD was measured every 15 min.

### Deep Mutational Scanning

Single site mutation libraries were generated by a PCR-based saturation mutagenesis method (*51*) as described below. The mutagenesis was carried out on the pUC19 plasmid containing the SH3 domain sequence of *ABP1* flanked by its homology arms subsequently used in the CRISPR mediated genomic re-integration. We used oligos containing degenerate nucleotides (NNN) to carry out the mutagenesis at each codon of the domain sequence. In the first step of the 2-step PCR procedure (short PCR step), an amplicon was generated with an oligo containing the degenerate codon which is positioned within the domain sequence, and another oligo lying outside Abp1 sequence in the plasmid. Short PCR step was carried out with the following settings: 5 min at 95°C, 35 cycles: 20 sec at 98°C, 15 sec at 60°C, 30 sec at 72°C and a final extension of 1 min at 72°C. The amplicon generated in this step was used as a megaprimer in the second step (long PCR step) to amplify the whole plasmid. Long PCR step was carried out with the following settings: 5 min at 95°C, 22 cycles: 20 sec at 98°C, 15 sec at 68°C, 3 min 30 sec at 72°C and a final extension of 5 min at 72°C. The long PCR product was digested for 2 hours at 37°C with DpnI. The digestion product was then transformed into *E.coli* MC1061 competent cells. In order to obtain most of the 64 possible codons, we recovered ~1000 colonies from each plate to retrieve the plasmid single mutant libraries. Saturation mutagenesis was carried out in a codon-position wise manner. Mutants at each position were stored separately. Library quality control was assessed by amplifying the purified plasmid DNA preparations by PCR followed by illumina sequencing as described below. The genomic insertion of the mutant libraries (one library per amino acid position) was performed by targeting the stuffer sequence in the *ABP1* SH3-deleted strain as described in the strain construction section. *ABP1* SH3-deleted yeast competent cells were co-transformed with pCAS containing the stuffer specific gRNA and PCR amplified *ABP1* SH3 single position mutated libraries from the pUC19 preparations. As for the library generation, ~1000 colonies from each plate were retrieved. Glycerol stocks of yeast cells were kept for each position (58 different stocks corresponding to the 58 amino acid positions of Abp1 SH3). The stocks were also validated by Illumina sequencing (see below).

### Liquid DHFR-PCA with librairies of mutants

Liquid DHFR-PCA experiments were performed following the generation of the *ABP1* SH3 single position mutated yeast strains and were performed in biological independent duplicates. An overnight culture was started in YPD + NAT by adding the same number of cells from each yeast mutant strains (master pool). Then, the master pool and Hua2-DHFR F[3] or Lsb3-DHFR F[3] preys strains were mixed in YPD (2:1) for mating and incubated for 8h at 30°C. Diploid cells were selected a first time by transferring the mixture in a YPD + NAT + HPH (OD 0.5) for 16h culture at 30°C. The next day, the first diploid selection was transferred in SC complet pH 6.0 + NAT + HPH at 0.5 OD and the culture was grown for 24h at 30°C. The validation of the yeast mutant libraries was done the next day with a fraction of the diploid cells (first DHFR-PCA experiment time point without PCA selection, reference condition S2) by amplifying *ABP1* SH3 genomic sequence followed by illumina sequencing as described in the section below. Following the selection, diploid cells optical density (OD) was read using a TECAN Infinite F200 Pro and dilutions to 0.1 OD in PCA selection media without agar in a 50 mL tube was done. Tubes were incubated at 30°C for 72 hours (second time point of the DHFR-PCA experiment, first PCA selection). A fraction of the cells (5U OD) were collected for illumina sequencing. The cells from the first PCA step were grown for a second PCA selection step using the same procedures. Finally, the genomic DNA of the cells (5U OD) after two cycles of PCA selection was extracted and the surviving *ABP1* SH3 mutant strains were detected by sequencing as described below.

### Single mutant libraries sequencing

For the validation of the plasmid single mutant libraries, the plasmid DNA was extracted from bacteria cells following transformation using a mini-prep plasmid extraction kit (FroggaBio). Yeast genomic DNA extraction of the library of *ABP1* SH3 mutants was performed using a standard phenol/chloroform protocol. Sequencing of yeast genomic DNA was performed at three different time points during the liquid DHFR-PCA experiments. After diploid selection of the bait-prey strains (single mutant library validation, reference condition S2) and following each of the two PCA selection rounds. For the final analysis presented in this article, the first and last time points were used. The libraries for sequencing were prepared by 3 successive rounds of PCR. The first PCR was performed with primers to amplify *ABP1* SH3 from saturation mutagenesis minipreps (4.5 ng of plasmid) or on genomic DNA extracted from yeast (90 ng of genomic DNA) (PCR program : 3 min at 98°C, 20 cycles : 30 sec at 98°C, 15 sec at 60°C and 30 sec at 72°C, final elongation 1 min at 72°C). The second PCR was performed to increase diversity in the libraries by adding Row and Column barcodes (*52*) for well identification in a 96 well plate (PCR program : 3 min at 98°C, 15 cycles : 30 sec at 98°C, 15 sec at 60°C and 30 sec at 72°C, final elongation 1 min at 72°C). The first PCR served as a template for the second one (2.25 μl of 1/2500 dilution). After the second PCR, 2 μl of the products were run on a 1.5% agarose gel and band intensity was estimated using Image Lab (BioRad Laboratories). The PCR products were mixed based on their intensity on agarose gel to roughly have equal amounts of each in the final library. Mixed PCRs were purified on magnetic beads and quantified using a NanoDrop (ThermoFisher). The third PCR was performed on 0.0045 ng of the purified pool from the second PCR to add a plate barcode and Illumina adapters (PCR program : 3 min at 98°C, 15 cycles : 30 sec at 98°C, 15 sec at 61°C and 35 sec at 72°C, final elongation 1 min 30 sec at 72°C). Each reaction for the third PCR was performed in triplicate, combined and purified on magnetic beads. After purification, libraries were quantified using a NanoDrop (ThermoFisher). Equal amounts of each library were combined and sent to the Genomic Analysis Platform (IBIS, Quebec, Canada) for pair-end 300bp sequencing on a MiSeq (Illumina).

### Analysis of DMS data

The raw data generated from deep sequencing (fastq format) was first demultiplexed based on the barcode sequences using custom scripts. Forward and reverse reads corresponding to each sample were merged using PEAR (*53*). The merged fastq reads were filtered to remove reads with average Phred score of less than 30 and the nucleotides with Phred score of less than 30 using fastp (*54*). The reads were aligned to their corresponding reference sequence using bowtie2 (*55*). Global alignment was carried out using the command: “bowtie2 -p 6 --end-to-end --very-sensitive --no-discordant --no-mixed -x $referencep -U $fastqp -S $samp”, where $referencep is path to the bowtie2 built reference, $fastqp is the path to the unaligned fastq file and $samp is the path to the aligned file in sam format. The aligned sam file was used to count the number of mutations using samtools (*56*) and pysam (https://github.com/pysam-developers/pysam). In order to normalise the counts by the depth of sequencing, the counts of the mutations were divided by the depth of sequencing at the position of the mutation. The normalised counts are referred to as frequencies. Next, the log2 ratios (pseudolog with pseudocount of 0.5) of frequencies in the test condition (second PCA selection) were normalized to the reference condition corresponding to the selection of diploids (before the PCA selections, S2) for each PPI. The python based code used for the analysis of the DMS data is available in the align module of ‘rohan’ package (*57*). Finally, the ratio of each mutation was scaled by subtracting the median ratio for synonymous codon substitutions (n=120) for each PPI (DMS score). Variants decreasing binding were defined based on the DMS score distribution for synonymous variants at the codon level. Mutations with a DMS score below the 1st percentile of this distribution represent ‘deleterious’ variants. Likewise, mutations improving PPIs have a DMS score above the 99th percentile of this distribution. For the structural analysis, the relative solvent accessibility (RSA) values were calculated by dividing surface accessibilities by the maximum surface accessibility for the relevant amino acid. Surface accessibilities for PDB: 2RPN (*22*) were calculated using the DSSP webserver (*58*), and the 20 maximum surface accessibility values (empirical) taken from (*59*). For the conservation analysis, the 27 yeast SH3s (excluding *SDC25-YLL017W*) were aligned using the MAFFT L-INS-i method (*60*), and then the per-site conservation calculated with a BLOSUM62 matrix using the *conserv(*) function of the R package ‘bio3d’.

### Cellular lysis and protein immunoprecipitation

Pre-cultures of yeast cells expressing baits fused with the DHFR F[1,2]-Flag were grown overnight at 30°C with agitation in synthetic complete medium. The next day, cells were expanded in synthetic complete medium to reach their exponential phase (OD of around 1.0), pelleted and frozen at −80°C. The equivalent of fifty OD600 of cells were resuspended in ice cold yeast lysis buffer (10 mM Tris-HCl pH 7.4, 150 mM NaCl, 0.5 mM EDTA (Millipore Sigma), 1% triton X-100 (Millipore Sigma) and one cOmplete Mini Protease Inhibitor Cocktail (Millipore Sigma) per 10 mL) with 425–600 M glass beads (Millipore Sigma) and disrupted at 4°C by 10 cycles of 2 min vortexing and 2 min cooldown. The equivalent of one hundred OD600 of lysed cells was used for the immunoprecipitation of the bait proteins using FLAG-M2 agarose beads (Millipore Sigma). The beads were incubated with the cell lysates for 120 mins at 4°C with slow rotation, followed by three washes with the yeast lysis buffer.

For the experiments with HEK293T cells (1×10cm confluent dish for WB and 3×15cm for MS), cells were washed in ice-cold PBS and lysed in lysis buffer (20 mM Tris-HCl pH 7.4, 150 mM NaCl, 1 mM EDTA, 0.5% sodium deoxycholate (Millipore Sigma), 10 mM beta-glycerophosphate (Millipore Sigma), 10 mM sodium pyrophosphate (Millipore Sigma), 50 mM NaF (Millipore Sigma)). Protease inhibitors were added to the lysis buffer (P8340 (Millipore Sigma) for WB and PMSF 1 mM, aprotinin 10 μg/mL, leupeptin 10 μg/mL and pepstatin 10 μg/mL (all Millipore Sigma) for MS). Cells were scraped in the lysis buffer and lysed at 4°C for 20 mins and centrifuged for another 20 mins at 20,000 g at 4°C. Bait proteins from the supernatant were immuno-precipitated for 90 mins at 4°C with FLAG-M2 agarose beads (Millipore Sigma) and washed three times in the lysis buffer. Beads were then used for WB or MS experiments (see below).

### Western blotting and antibodies

Protein extracts and affinity-purified baits from yeast and human lysed cells were migrated on 10% polyacrylamide gels. The separated proteins were then transferred on nitrocellulose membranes. Validation of correct protein loading was done with ponceau staining of the membranes. Before the primary antibody incubations, membranes were blocked in 5% milk. Protein signals were detected on an Amersham Imager 600RGB (GE Healthcare) following the incubation of the membranes with the appropriate HRP-conjugated secondary antibodies and the Clarity Western ECL Substrate (Bio-Rad). The antibodies used in this study are: M2-HRP (A8592, Sigma), PAK (SC-881, Santa Cruz), p130cas/BCAR1 (SC-860, Santa Cruz), actin (Cell Signaling Technology), anti-mouse HRP (Cell Signaling Technology) and anti-rabbit HRP (Cell Signaling Technology). The quantification of the WB signals was performed with the Amersham Imager 600 analysis software (GE Healthcare).

### Live imaging of endocytosis

Overnight cultures of Sla1-GFP cells were diluted to an OD600 of 0.15 and grown in synthetic complete medium without tryptophan until they reached an OD600 of ~0.4-0.5 at 30°C. The cells were then seeded on concanavalin A 0.05 mg/mL (Millipore Sigma) coated 8-well glass-bottom chamber slides (Sarstedt). Image acquisition was performed using a Perkin Elmer UltraVIEW confocal spinning disk unit attached to a Nikon Eclipse TE2000-U inverted microscope equipped with a Plan Apochromat DIC H 100×/1.4 oil objective (Nikon), and a Hamamatsu Orca Flash 4.0 LT+ camera. Imaging was done at 25°C in an environmental chamber. The software NIS-Elements (Nikon) was used for image capture. For each field, one brightfield and a series of fluorescence (GFP) images were taken. Cells were excited with a 488 nm laser and emission was filtered with a 530/630 nm filter. GFP time laps were acquired continuously at a rate of 1 frame/sec for 3 minutes.

### Analysis of microscopy images

Bright field images were used to segment cells using YeastSpotter (*61*). Cells were filtered based on circularity, solidity and the normalized difference between minor and major axis lengths to remove poorly detected cells. Out of focus cells were also filtered out based on size and brightness. The centroids of the segmented cells were used to identify the locations of each cell in the image using Scikit-Image (*62*). The location of each cell was used to isolate individual cell time-frames of fluorescence (GFP) images. Each cell time frame of the fluorescence images was processed through a python based single particle tracking tool - Trackpy (*63*). Trackpy detected the locations of Sla1-GFP positive foci in each frame and linked them together to provide particle-wise trajectories. Trajectories detected in less than 10 frames were considered as spurious and were filtered out. Segments of the trajectories spuriously indicating movement of the particles back to the cell membrane were trimmed. The preprocessed trajectories of the particles were then used for the calculation of distances using the spatial distance module of ‘SciPy’ (*64*). The python based code used for the analysis of microscopy images are included in the endocytosis module of ‘htsimaging’ package (*65*).

### Protein purifications

*E. coli* BL21 (DE3) cells were transformed with mNCK2 constructs in the pET-30b vector and grown overnight at 37°C in LB with 100 μg/ml kanamycin. The next day, cultures were expanded in LB with kanamycin until they reached their exponential growth phase (OD of around 1.0). Protein production was then induced by the addition of 1 mM IPTG (Bio Basic) and incubation at 16°C overnight with agitation. Cultures were pelleted with a centrifugation step (3,200 g at 4°C, 30 mins), washed with ice-cold PBS and pellets were kept at −80°C. Bacteria were thawed in GST buffer (20 mM Tris-HCl pH 7.5, 0.5 M NaCl, 1 mM EDTA and 1 mM DTT (Millipore Sigma)) with cOmplete protease inhibitors (Millipore Sigma) and disrupted with sonication. Lysates were then centrifuged at 20,000 g for 20 min at 4°C and the supernatant loaded to a GSTrap FF 1 mL column (GE Healthcare). Following a washing step with 10 mL GST-Buffer, the GST-Tagged proteins were eluted with 15 mM GSH (Bio-basic) and cleaved in solution using purified recombinant His-Tagged TEV protease (Thermo Fisher Scientific). Tag-free NCK2 proteins were further purified using a 1 ml HisTrap FF column (GE Healthcare) to remove free His-tagged GST and TEV as well as remaining impurities. The purified proteins were concentrated with Amicon 10K filters (Millipore Sigma) and their concentration assessed using Coomassie and BCA assays (Thermo Fisher Scientific).

### Fluorescence polarization

Increasing amounts of NCK2 full-length recombinant proteins in GST buffer (final concentrations of 0-150 μM for CD3ε and 0-175 μM for PAK1) were used for binding assays with a constant amount of fluorescein isothiocyanate (FITC)-conjugated peptides (resuspended in 17 mM Tris-HCl pH 8, 100 mM NaCl and 0.5% Brij L23 (Millipore Sigma), peptides final concentration of 10 nM). Cd3ε (RGQNKERPPPVPNPDY) and PAK1 (DIQDKPPAPPMRNTST) peptides were used as NCK2 SH3-1 and NCK2 SH3-2 ligands, respectively. Binding assays were performed in the FP buffer (17 mM Tris-HCl pH 8 and 100 mM NaCl) and fluorescence polarization was measured on a Synergy H1 multi-mode plate reader (Bio Tek) at 535 nm with 485 nm excitation. Calculation of the dissociation constants was performed with a one-site total binding model and nonlinear regression in GraphPad Prism version 7.

### Phase separation

Purified NCK2 proteins were diluted in GST buffer to identical concentrations. At 500 mM NaCl present in this buffer, all NCK2 constructs were completely soluble. Phase transition was initiated by dilution in imidazole buffer (10 mM imidazole (Bio Basic) pH 7.0, 1 mM DTT, 1mM EGTA (Millipore Sigma), 1 mM MgCl2, 2.5% Glycerol (Bioshop Canada), 3% Dextran (Millipore Sigma)) containing various amount of NaCl so that the final concentration of NaCl would correspond to those in Fig. S7B. For all the other experiments, phase transition of NCK2 proteins was assessed at 60 mM NaCl. NCK2 at 8 μg/μL (40 μg in 5uL) was added to 35 uL imidazole buffer in a plastic 96 well plate that had first been pre-treated with a 0.2% BSA solution to reduce nonspecific binding. The plate assay was then incubated overnight (24 hours) at 4°C before imaging. Phase contrast images of each well were acquired using a Zeiss Axio Vert.A1 inverted microscope with a LD A-Plan 40x/0.55 Ph1 objective (Zeiss) in Zen Blue software (version 2.3.69.1000). After imaging, the soluble protein fraction (supernatant) was recovered for SDS-PAGE analysis. Protein droplets attached at the bottom of the well (pellet) were washed once with imidazole buffer with 60 mM NaCl and then solubilized in Laemmli buffer. The pellets to supernatant fractions were used to quantify phase transition at 60 mM NaCl (7 replicates) via Coomassie protein quantification. The involvement of weak hydrophobic interactions in NCK2 driven phase transition was assessed using 1,6-Hexanediol (Millipore Sigma) at 60 mM NaCl. Imidazole buffer containing 0, 1%, 5% and 10% 1,6-Hexanediol was first prepared and phase transition was initiated and analyzed as described above. Finally, competition of SH3-2 dependent homotypic NCK2 interaction with a SH3-2 binding peptide was assessed in the presence of 5 μM, 15 μM, 45 μM, 90 μM, 150 μM, or 300 μM PAK1 peptide (DIQDKPPAPPMRNTST) in imidazole buffer with 60 mM NaCl. The impact of PAK1 peptide addition was observed via phase contrast microscopy as described above.

### Experimental design for mass spectrometry experiments

NCK2 baits purified on beads from HEK293T cells (as previously described) were washed two additional times in 20 mM Tris-HCl pH 7.4 and eluted in 50 mM phosphoric acid. Proteins were then digested with trypsin (Promega) on-beads as previously detailed (*66*). Briefly, eluted proteins were concentrated on ZipTip^®^-SCX (Millipore Sigma), reduced with TCEP 100 mM (Millipore Sigma) and alkylated during the tryptic digestion with iodoacetamide 10 mM (Millipore Sigma). Peptides were then eluted in ammonium bicarbonate 200 mM (Millipore Sigma) and desalted on Stage Tips C18 columns (*67*). The final eluted peptides were dried using an Eppendorf Vacufuge™ Concentrator. For each bait, two biological replicates were processed independently. These were analyzed alongside four negative controls corresponding to transiently expressed 3×FLAG-GFP purified from HEK293T cells (as described for NCK2 constructs). These control cell lines were grown in parallel to those expressing baits and treated in the same manner. NCK2 W38/W148/W234K triple mutant, with the tryptophan residue essential for SH3 interactions mutated in each SH3, was used as a SH3-inactive negative control for SH3-dependent interactions. The equivalent of one third of each sample was used for the SWATH-libraries generation and another third for SWATH quantification. Data dependent acquisition (DDA) and data independent acquisition (DIA) analysis of each sample was performed back-to-back without washes to diminish the instrument time required to complete the analysis. To minimize carry-over issues during liquid chromatography, extensive washes were performed between each sample; and the order of sample acquisition on the mass spectrometer was also reversed for the second biological replicates to avoid systematic bias. We validated that the same quantity of baits was used for each NCK2 chimeras by quantifying NCK2 peptides in each sample (see Table S6).

### Proteins identification by mass spectrometry

The analyses were performed at the proteomic platform of the Quebec Genomics Center. Peptide samples were separated by online reversed-phase (RP) nanoscale capillary liquid chromatography (nanoLC) and analyzed by electrospray mass spectrometry (ESI MS/MS). The experiments were performed with a Dionex UltiMate 3000 nanoRSLC chromatography system (Thermo Fisher Scientific) connected to a TripleTOF 5600+ mass spectrometer (Sciex) equipped with a nanoelectrospray ion source. Peptides were trapped at 20 μl/min in loading solvent (2% acetonitrile, 0.05% TFA) on a 5 mm x 300 μm C_18_ pepmap cartridge pre-column (Thermo Fisher Scientific) during 5 minutes. Then, the pre-column was switch online with a self-made 50 cm x 75 um internal diameter separation column packed with ReproSil-Pur C_18_-AQ 3-μm resin (Dr. Maisch HPLC) and the peptides were eluted with a linear gradient from 5-40% solvent B (A: 0,1% formic acid, B: 80% acetonitrile, 0.1% formic acid) in 120 minutes, at 300 nL/min. In DDA, the instrument method consisted of one 250 milliseconds (ms) MS1 TOF survey scan from 400–1300 Da followed by 20 100 ms MS2 candidate ion scans from 100–2000 Da in high sensitivity mode. Only ions with a charge of 2+ to 5+ that exceeded a threshold of 150 cps were selected for MS2, and former precursors were excluded for 20 s after one occurrence. In DIA, acquisition consisted of one 50 ms MS1 scan followed by 32 × 25 a.m.u. isolation windows covering the mass range of 400–1250 a.m.u. (cycle time of 3.25 s); an overlap of 1 Da between SWATH was preselected. The collision energy for each window was set independently as defined by CE = 0.06 × m/z + 4, where m/z is the centre of each window, with a spread of 15 eV performed linearly across the accumulation time.

### Data Dependent Acquisition (DDA) MS analysis

Mass spectrometry data was stored, searched and analyzed using the ProHits laboratory information management system (LIMS) platform (*68*). Sciex .wiff mass spectrometry files were converted to mzML and mzXML using ProteoWizard (3.0.4468;(*69*)). The mzML and mzXML files were then searched using Mascot (v2.3.02) and Comet (v2012.02 rev.0) (*70*). The spectra were searched with the RefSeq database (version 57, January 30th, 2013) acquired from NCBI against a total of 72,482 human and adenovirus sequences supplemented with “common contaminants” from the Max Planck Institute (http://141.61.102.106:8080/share.cgi?ssid=0f2gfuB) and the Global Proteome Machine (GPM; http://www.thegpm.org/crap/index.html) (*71*). Charges +2, +3 and +4 were allowed and the parent mass tolerance was set at 12 ppm while the fragment bin tolerance was set at 0.6 amu. Carbamidomethylation of cysteine was set as a fixed modification. Deamidated asparagine and glutamine and oxidized methionine were allowed as variable modifications. The results from each search engine were analyzed through TPP (the Trans-Proteomic Pipeline (v4.6 OCCUPY rev 3), (*72*) via the iProphet pipeline (*73*)).

### Data independent acquisition (DIA) MS analysis with MSPLIT-DIA

DIA MS data was analysed using MSPLIT-DIA (version 1.0; (*74*)) implemented in ProHits 4.0 (*68*). To generate a sample-specific spectral library for the FLAG AP-MS dataset, peptide-spectrum matches (PSMs) from matched DDA runs (18 runs) were pooled by retaining only the spectrum with the lowest MS-GFDB (Beta version 1.0072 (6/30/2014) (*75*)) probability for each unique peptide sequence and precursor charge state, and a peptide-level false discovery rate (FDR) of 1% was enforced using TDA (*76*). The MS-GFDB parameters were set to search for tryptic cleavages, allowing no missed cleavage sites, 1 C_13_ atom per peptide with a mass tolerance of 50 ppm for precursors with charges of 2+ – 4+ and a tolerance of ±50 ppm for fragment ions. Peptide length was limited to 8–30 amino acids. Variable modifications were deamidated asparagine and glutamine and oxidized methionine. The spectra were searched with the NCBI RefSeq database (version 57, January 30th, 2013) against a total of 36,241 human and adenovirus sequences supplemented with “common contaminants” from the Max Planck Institute (http://141.61.102.106:8080/share.cgi?ssid=0f2gfuB) and the Global Proteome Machine (GPM; http://www.thegpm.org/crap/index.html). Decoys were appended using the decoy library command built in to MSPLIT-DIA, with a fragment mass tolerance of ±0.05 Da. The spectral library was then used for protein identification by MSPLIT as previously described (*74*) with peptides identified by MSPLIT-DIA passing a 1% FDR subsequently matched to genes using ProHits 4.0 (*68*). The MSPLIT search parameters were as follows: parent mass tolerance of ±25 Da and fragment mass tolerance of ±50 ppm. When retention time was available within the spectral library, a cut-off of ±5 min was applied to spectral matching as previously described (*74*).

### MS data analysis with SAINTexpress

SAINTexpress version 3.6.1 (*77*) was used as a statistical tool to calculate the probability value of each potential protein-protein interaction compared to background contaminants using default parameters. The four control samples were used in uncompressed mode. Two unique peptide ions and a minimum iProphet probability of 0.95 were required for protein identification prior to SAINTexpress in DDA mode while only 2 unique peptides were required in DIA mode.

### Position and sequence conservation of orthologs

The UniProt database was used to retrieve full-length sequences of all *S. cerevisiae* SH3 proteins except for the Sdc25 (YLL017W) pseudogene (*78*). One-to-one orthologs for each protein were then retrieved from fungal species using Ensembl Compara (*79*). This process was automated for each SH3 protein using the Ensembl REST API (*80*). The *hmmscan* function of HMMER v3.3 was then used for the domain annotation of all orthologs using default significance thresholds of 0.01 (E = 0.01 and domE = 0.01) and the Pfam-A HMM library (*81*). Any domain belonging to the Pfam SH3 clan (CL0010) was assigned as an SH3 domain. SH3 domain positions for each orthologous sequence were taken by dividing the domain start point (‘env_from’) by the full sequence length. This calculation was repeated for each *S. cerevisiae* homolog, and orthologous SH3 domains were considered positionally conserved if their domain positions were within a 10% sequence length window of the corresponding *S. cerevisiae* domain.

The sequence conservation of each SH3 domain was calculated using the same set of orthologous sequences as described directly above. Domain sequences within an orthologous group were aligned together using the MAFFT L-INS-i method (*60*). The pairwise similarity between domain sequences was determined from the *seqidentity(*) function in the R package ‘bio3d’ (*82*), and we took the mean pairwise sequence similarity between the *S. cerevisiae* sequence and all its orthologs as a measure of sequence conservation for each domain.

### Tree comparison analysis

In Figure 2C, the relationship between SH3 domain sequences and their PPI profiles in Abp1 was explored by constructing dendrograms for both features and then quantifying the similarity between dendrograms. For the interaction similarities (Figure 2C, left), a Euclidean distance matrix of the PCA scores was first constructed using the *dist(*) function in R, and then the distance matrix used to generate a dendrogram from the *hclust(*) function in R with default parameters. For the sequence similarities (Figure 2C, right), the SH3 domain sequences were first aligned using the MAFFT L-INS-i method (*60*). A distance matrix was then constructed from the *seqidentity(*) function in the R package ‘bio3d’ (*82*) — by taking *distance* = *1-similarity* — and then hierarchical clustering performed in the same manner as for the DHFR-PCA PPIs. Side-by-side visualisations of the dendrograms were generated using the R package ‘dendextend’ (*83*). The overall similarity between dendrograms was quantified using the cophenetic correlation, which correlates the tree-wise distances for both dendrograms across all possible SH3 pairs. This was calculated using the *cor.dendlist(*) function in the ‘dendextend’ package. To test for statistical significance, we permuted the data by randomising the assignment of SH3 domains to different DHFR-PCA PPIs profiles and then recalculated the cophenetic correlation using all steps described directly above. This was repeated 10,000 times to generate a random distribution of cophenetic correlation scores. Finally, a similar tree-based approach was used to compare PPI profiles with growth profiles for Sla1 constructs generated from the domain shuffling experiments (Fig. S5B). Both the DHFR-PCA interaction data and the Sla1 growth data was clustered as described above, by generating a Euclidean distance matrix and then performing hierarchical clustering.

### PWMs enrichment analysis

All yeast PWMs were taken from a 2009 study of SH3 domain specificity (*12*). The similarity between SH3 PWMs and known interactors was assessed using a matrix similarity score (MSS) derived from the MATCH algorithm (*84*). This scoring method assigns a score of 1 to perfect sequence matches to the PWM and vice versa. For each interactor assigned to an SH3 domain, MSS scores for the corresponding PWM were calculated for all possible sequence *k*-mers (*k* = number of PWM columns) and then the maximum taken (Max. MSS). This procedure was repeated for all SH3 domains with an assigned PWM and then the results pooled to generate Figures 1C, 1D, and Figure S2D. Some SH3 domains were represented by more than one PWM from (*12*), reflecting multiple specificities (*85*). In these cases, MSSs were calculated for both PWMs and then the overall maximum score taken forward for further analysis.

In Figure 1, maximum MSS scores were calculated for sequences belonging to the ‘Random’, ‘SH3-independent PPI’, ‘SH3-inhibited PPI’, and ‘SH3-dependent PPI’ category of interactor sequences. For the ‘Random’ category, random peptides were generated by sampling amino acids according to their background frequency in the *S. cerevisiae* proteome. For SH3-independent PPI, we included only those interactors that were found to be unaffected by *any* SH3 domain deletions given that *in vitro-derived* SH3 PWMs overlap strongly, which could lead to spurious MSS enrichment for SH3-independent PPIs. In Figure 1D, maximum MSSs were also re-calculated for the ‘SH3-dependent PPI’ category of interactor sequences after randomly assigning PWMs to each SH3 domain; this procedure was repeated until 10,000 maximum MSS scores were sampled. For Figure S2D, the ‘PPI gained by Abp1’ correspond to PPIs gained by Abp1 after domain swapping, and the ‘PPI unaffected’ correspond to PPIs of Abp1 that do not change after domain swapping. The ‘Random’ category was generated using the same approach described directly above for Figure 1.

### Comparison of interactome with literature

To find previously reported PPIs, we parsed a recent release (v 3.5.16) of the BioGRID and searched for reports of physical interactions between baits and preys (*11*). PPIs were searched in the two directions possible (A-B and B-A) in the database.

### GO enrichment

To determine the biological processes enriched in our DHFR-PCA data, our list of positive interactions partners were analyzed against our list of unique preys using the Gene Ontology Term Finder of the SGD database, only considering enrichment with p-value <= 0.01 (*86*).

## QUANTIFICATION AND STATISTICAL ANALYSIS

All the statistical details of the different experiments can be found in the figure legends, figures and in the results sections.

---

**Fig. S1.**
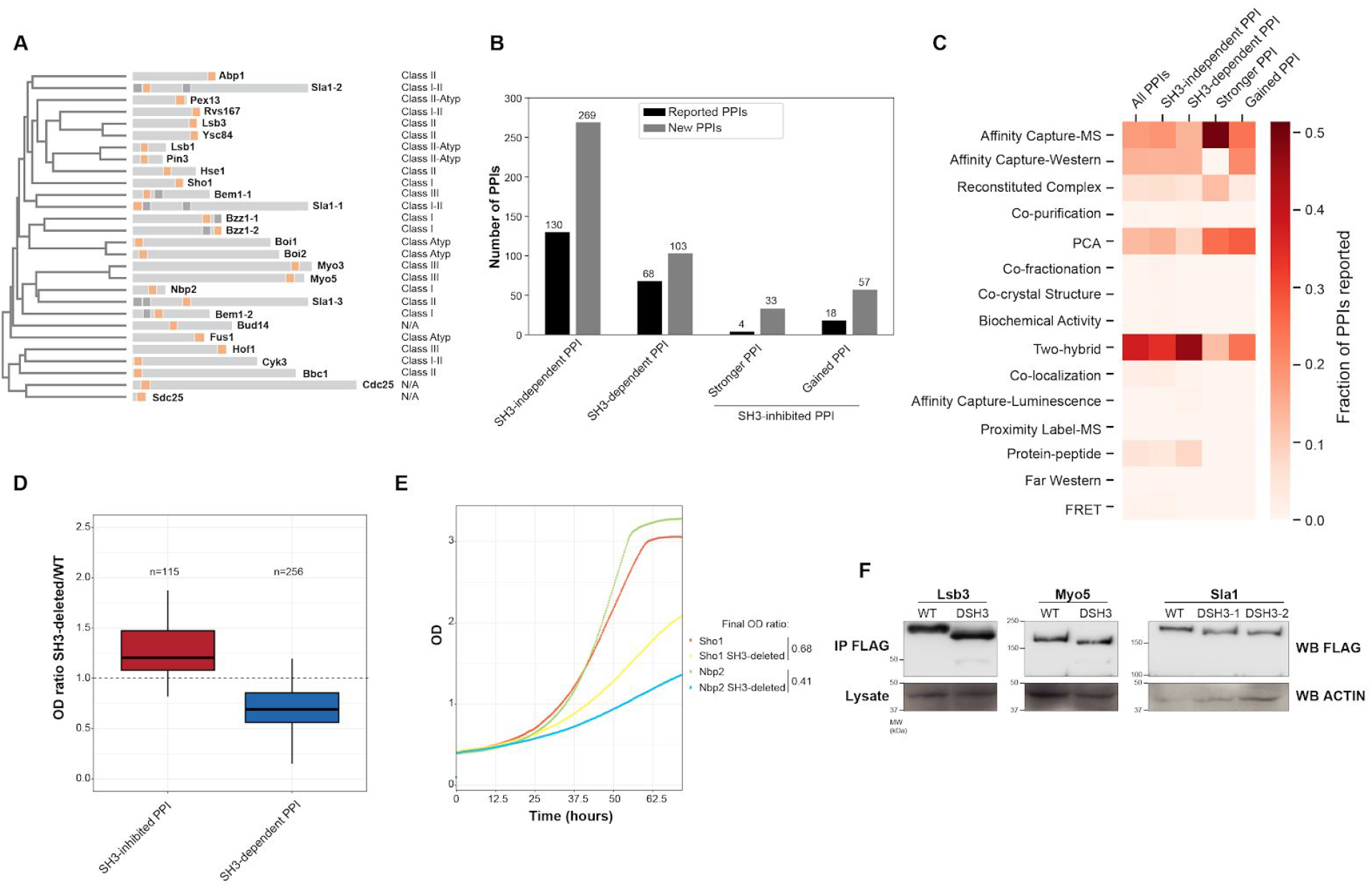
Yeast SH3 domains, the number and types of PPI changes in response to SH3 deletion and validation. A) The relationships of yeast SH3 amino acid sequences. Domain position and length relative to their host proteins are illustrated. SH3s used in the phylogeny are shown in orange. Grey squares are other SH3s in the same protein. The preferred type of binding motifs from *in vitro* assays are indicated (*12*). B) The proportion of PPIs previously reported for the different types of PPIs affected or not by SH3 deletion (*11*). C) The fraction of known PPIs per method of detection is shown for the same categories of interactions as in B) (*11*). D) Validation of the PPIs altered by the deletion of yeast SH3s using low-throughput liquid DHFR-PCA. The ratios (SH3-deleted/WT) of the optical density from the last time point of the experiment for the growth curves of each PPI are shown. A ratio higher than 1 represents an interaction that is stronger upon SH3 deletion. E) Examples of two SH3-dependent PPIs DHFR-PCA liquid assay. Growth curves are shown for the WT and SH3-deleted baits for Sho1-Pbs2 and Nbp2-Pbs2 PPIs. The ratios of the final optical density (SH3-deleted/WT) as represented in panel D) are also indicated. F) Western blot analysis of the expression level of SH3-deleted proteins that lost many PPIs upon their SH3-deletion. All baits have a C-terminal 1xFLAG tag allowing their immuno-detection. See also Table S1 and S2.

**Fig. S2.**
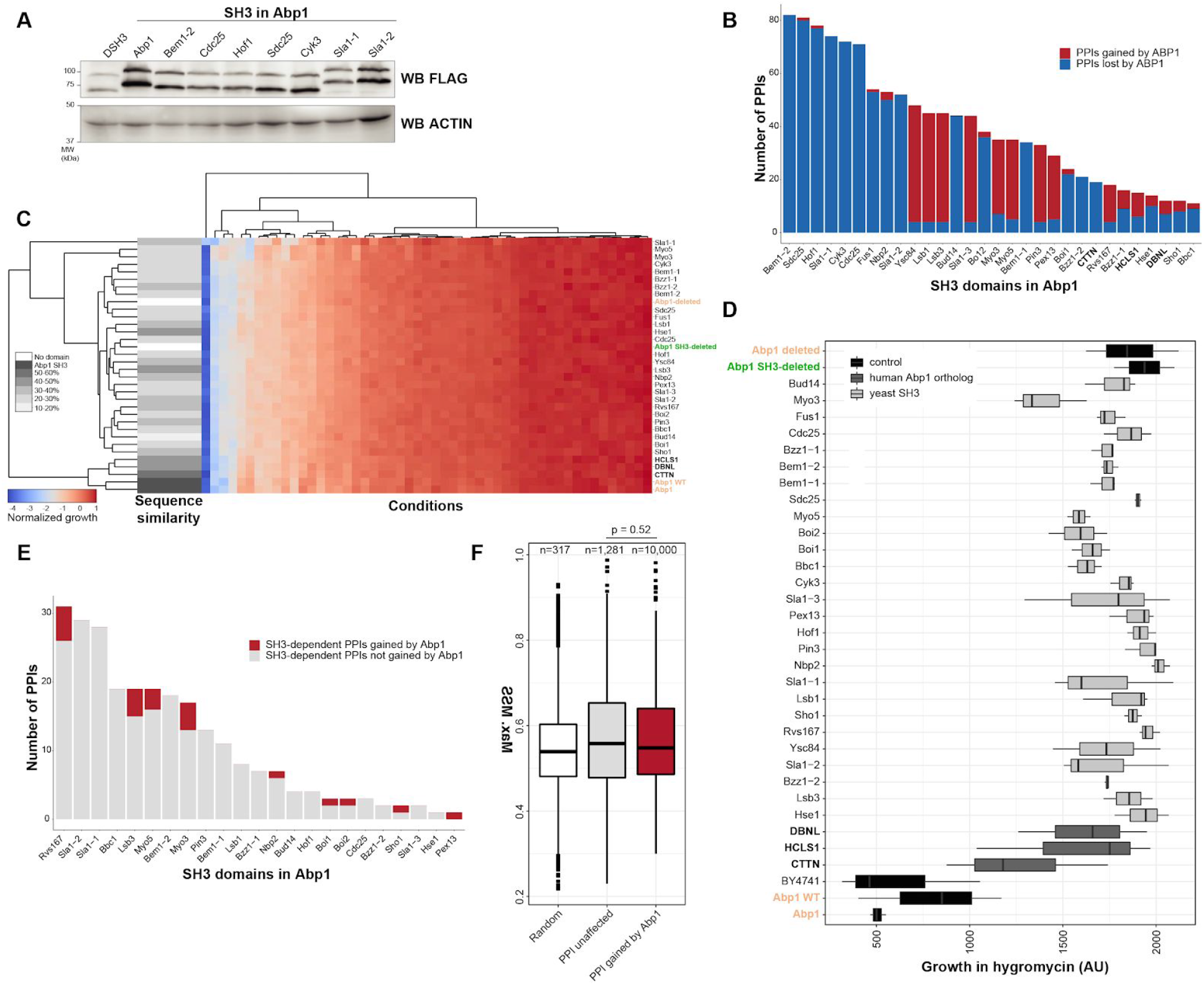
Characterization of Abp1 SH3 domain swapping. A) Protein expression levels of the Abp1 SH3 swapped proteins for which the smallest number of DHFR-PCA PPIs were measured as compared to the Abp1_SH3_ in Abp1 and SH3-deleted strains. Two bands are detected for Abp1, as previously observed (*41*). B) Number of PPIs that were affected by Abp1 SH3 domain swapping. C) Growth in stress conditions for the Abp1 SH3 swapped strains. The growth values were scaled per strain (row). Blue to red represents the normalized growth per strain after 74 hours (log2 colony size). The sequence similarity of each SH3 to Abp1 SH3 is represented in grey scale. Each strain was grown in twelve replicates. D) Growth of Abp1 SH3 swapped stains in liquid medium with hygromycin is represented (area under the curve (AU)). The control strains, shown in black, highlight that the resistance to the drug is dependent on Abp1 SH3. Domains are sorted by increasing sequence similarity with Abp1 from the top. Growth rates were measured in triplicates. E) Number of SH3-dependent PPIs that were gained by each SH3 swapped to Abp1 (as determined in Figure 1B) in comparison to the ones not gained. F) PWMs analysis of PPIs gained by Abp1 using the matrix similarity scores (MSS) between the PWM and sequence. This analysis shows no enrichment of the predicted SH3 motifs in the gained PPIs relative to the unaffected Abp1 PPIs (p = 0.52, Mann-Whitney test, one-sided). For C) and D) the orange SH3s are controls and the SH3-deleted protein is in green. In all panels, human SH3s are shown in bold. See also Table S3.

**Fig. S3.**
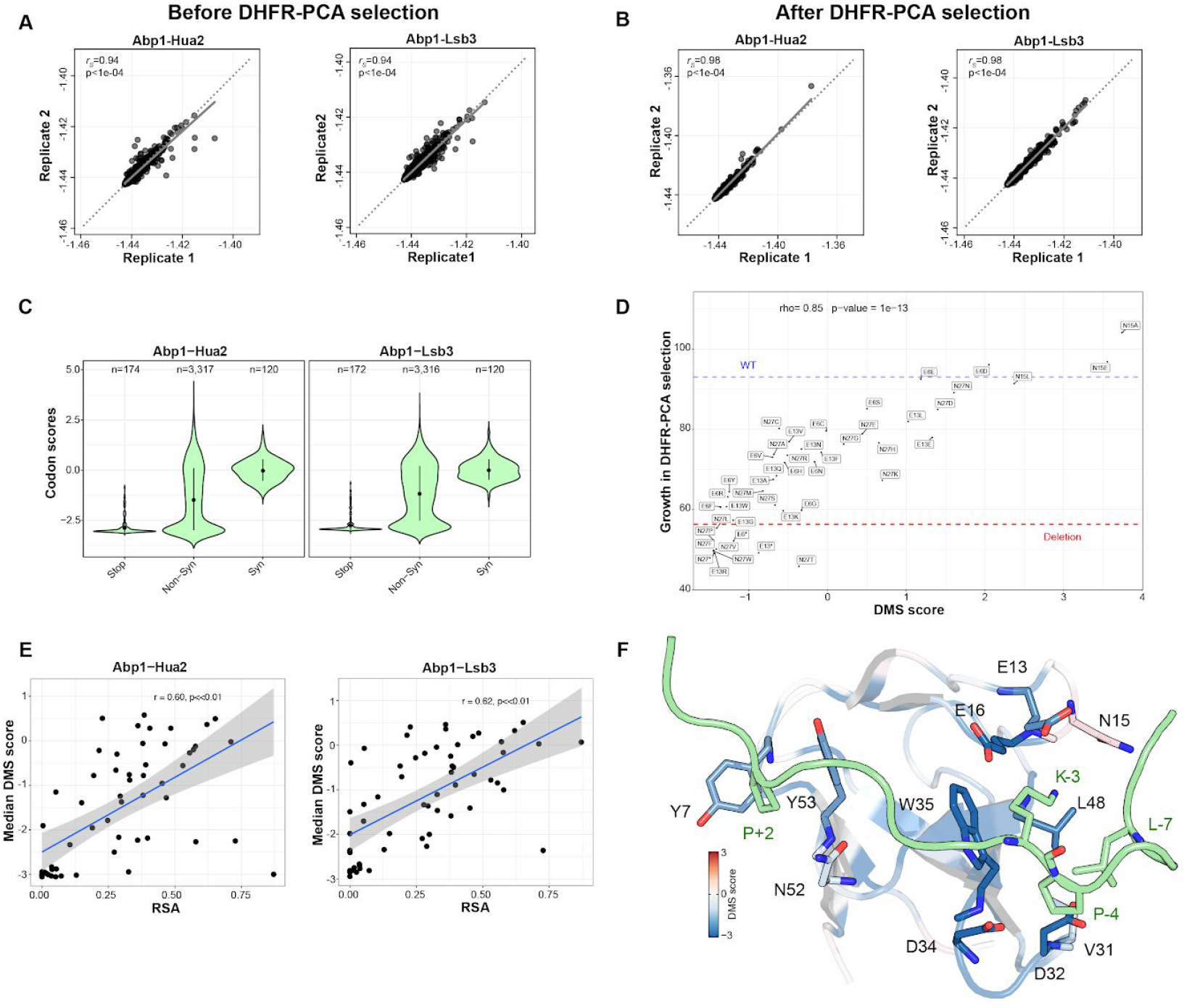
Characterization and validation of DMS mutants of Abp1 SH3 domain. A-B) Reproducibility between biological replicates of DMS DHFR-PCA experiments. The correlations for the two interactors are shown (A is for the reference condition and B for the DHFR-PCA condition). The frequencies of mutants (shown on log2 scale) were compared in terms of Spearman’s rank correlation coefficient (r_s_). Associated p-value is shown on each plot. C) Distribution of the codon log2 average sequence counts for each type of mutation of Abp1 SH3 for the two PPIs. Syn is for synonymous. D) Selected non-conserved positions (E6, E13, N15 and N27) sensitivities to mutations for the Abp1-Hua2 PPI were tested in a low-throughput liquid DHFR-PCA assay. The growth of the mutants in the low-throughput experiment (average of four replicates of the total OD for each growth curve, y axis) is compared to the mutants DMS score (x axis). The growth of WT Abp1 and Abp1 SH3-deleted are represented by horizontal dash lines. E) The median DMS score of the 58 different Abp1 SH3 positions for both PPIs in relation to their relative solvent accessibility (RSA). F) Abp1 SH3 residues in close proximity (within 4 Å) to the most important binding residues of the Ark1 peptide (P+2, K-3, P-4, and L-7) are mapped on the structure of Abp1 SH3 in complex with the peptide (PDB: 2RPN, (*22*)). Ark1 peptide is shown in green, oxygen atoms are in red and nitrogen atoms are in blue. The colors of Abp1 SH3 residues that are mapped on the structure represent their average DMS score for Abp1-Hua2 PPI. Most interface residues sensitive to mutations, such as Y7, E13, E16, V31, D32, D34, L48 and Y53, specifically affect Hua2 PPI. See also Table S4.

**Fig. S4.**
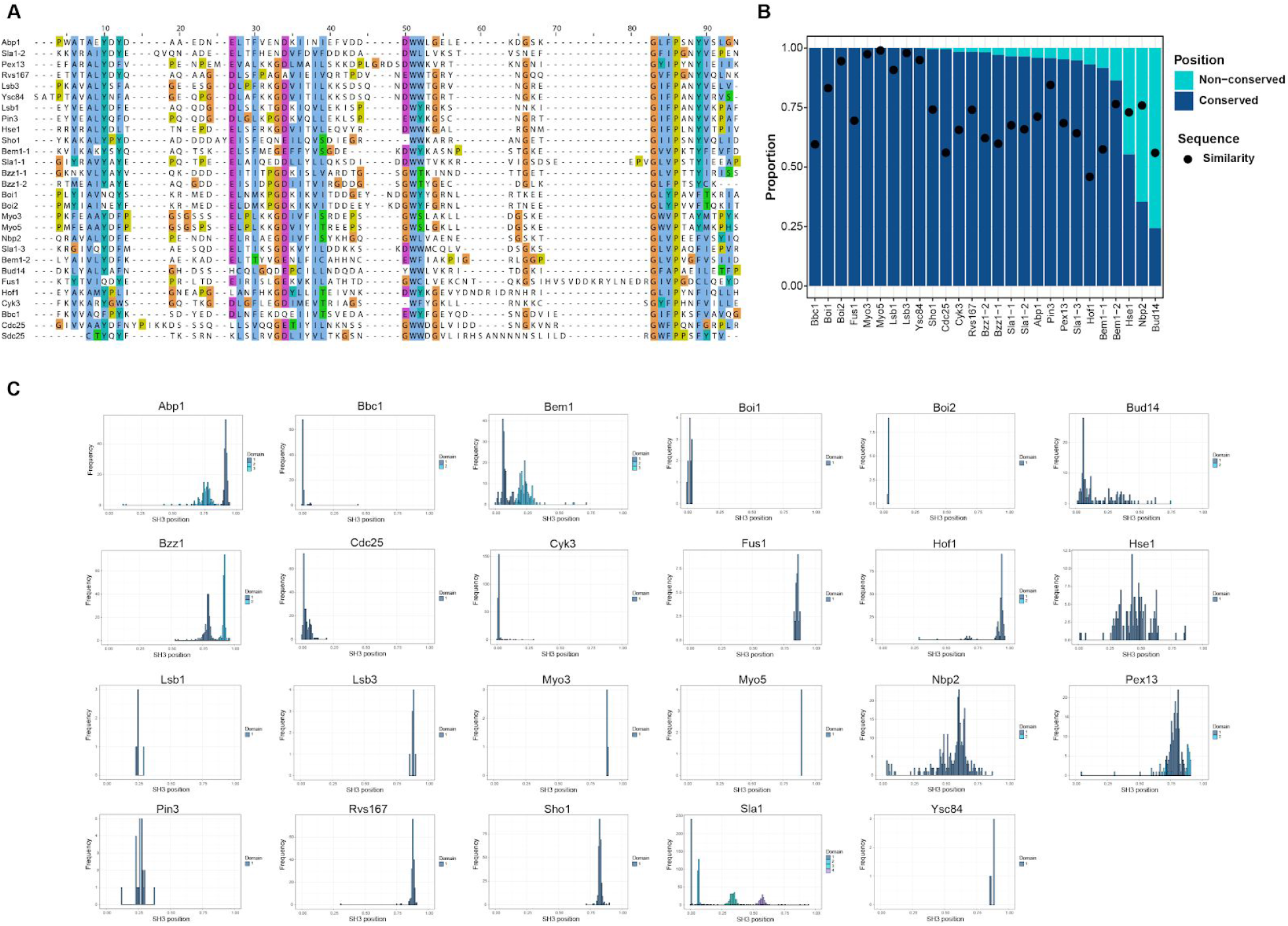
Yeast SH3 domain sequence and position in their host protein: A) Multiple sequence alignment of yeast SH3 domains. B) The y-axis shows the fraction of one-to-one orthologs (across 449 fungal species) with conserved domain positions relative to *S. cerevisiae*. The mean sequence similarity is represented by black circles. The pseudogene Scd25 was excluded from these analyses. C) Each panel represents the distribution of SH3 domain positions for the one-to-one orthologs of a given *S. cerevisiae* SH3. For each ortholog sequence, SH3 domain start positions were identified from SH3 hidden Markov models (HMMs), and then the start positions compiled across all orthologs to derive a frequency distribution. The SH3 domains are numbered in cases where more than one SH3 was identified across orthologs, where ‘1’ represents the domain that is also found in the *S. cerevisiae* copy. For multi-SH3 proteins (Bem1, Bzz1, and Sla1) the numbers correspond to the domain numbering given in Fig. S1A. See also Table S1.

**Fig. S5.**
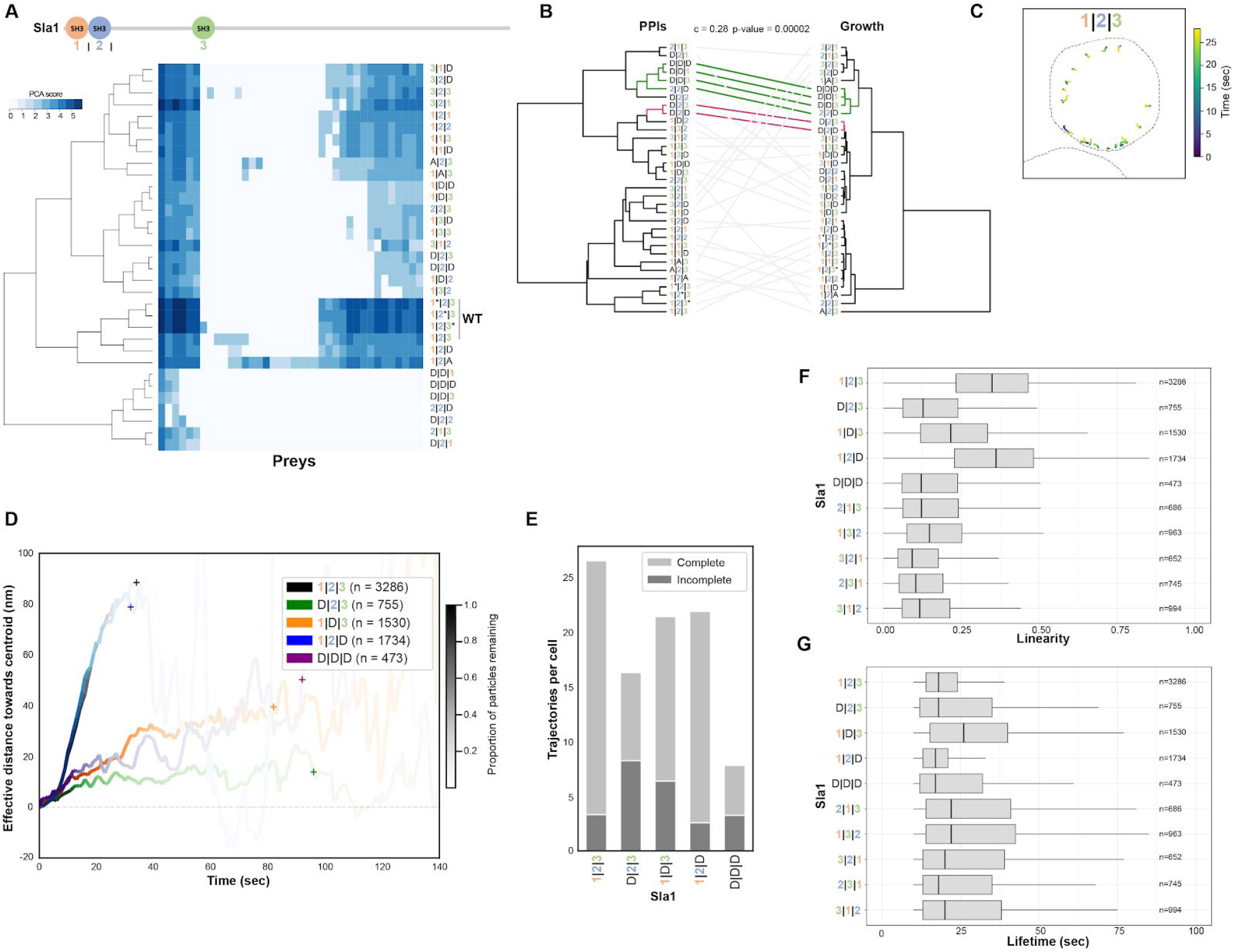
SH3 shuffling impacts Sla1 PPIs, cells growth in stress conditions and clathrin-mediated endocytosis. A) Sla1 SH3-deleted or -shuffled PPIs as detected by DHFR-PCA. The color code represents PPI strength as detected by DHFR-PCA (PCA score). All PPIs were measured in quadruplicate. A is for Abp1 SH3 and * is for the reinsertion of the WT domain as control strains. A scaled cartoon of Sla1 is also illustrated. B) Cophenetic correlation for the similarity of Sla1 SH3-shuffled PPI clusters with the growth phenotype clusters. Empirical p-value obtained from data permutation is p = 0.00002. C) Representative fluorescence microscopy timeframe analysis of a yeast cell expressing WT Sla1-GFP. The calculated trajectories for every Sla1-GFP foci detected are shown. Cell membrane is delimited with a dashed line and each Sla1-GFP particle colors represent its position in time. D) Sla1-GFP particles average effective distance travelled towards the cell center through time. Sla1 SH3-deleted strains are shown. The proportion of events that are not completed yet is represented by the color transparency of the curves (+ represent the time point when 95% of foci have disassembled). E) Complete or incomplete Sla1-GFP endocytosis events are shown (average per cell) for Sla1 SH3-deleted strains. F) Linearity of Sla1-GFP particle trajectories for each SH3 deletion or shuffling. G) Sla1-GFP particle lifetime (in seconds) are shown for the same strains as in F). See also Table S5.

**Fig. S6.**
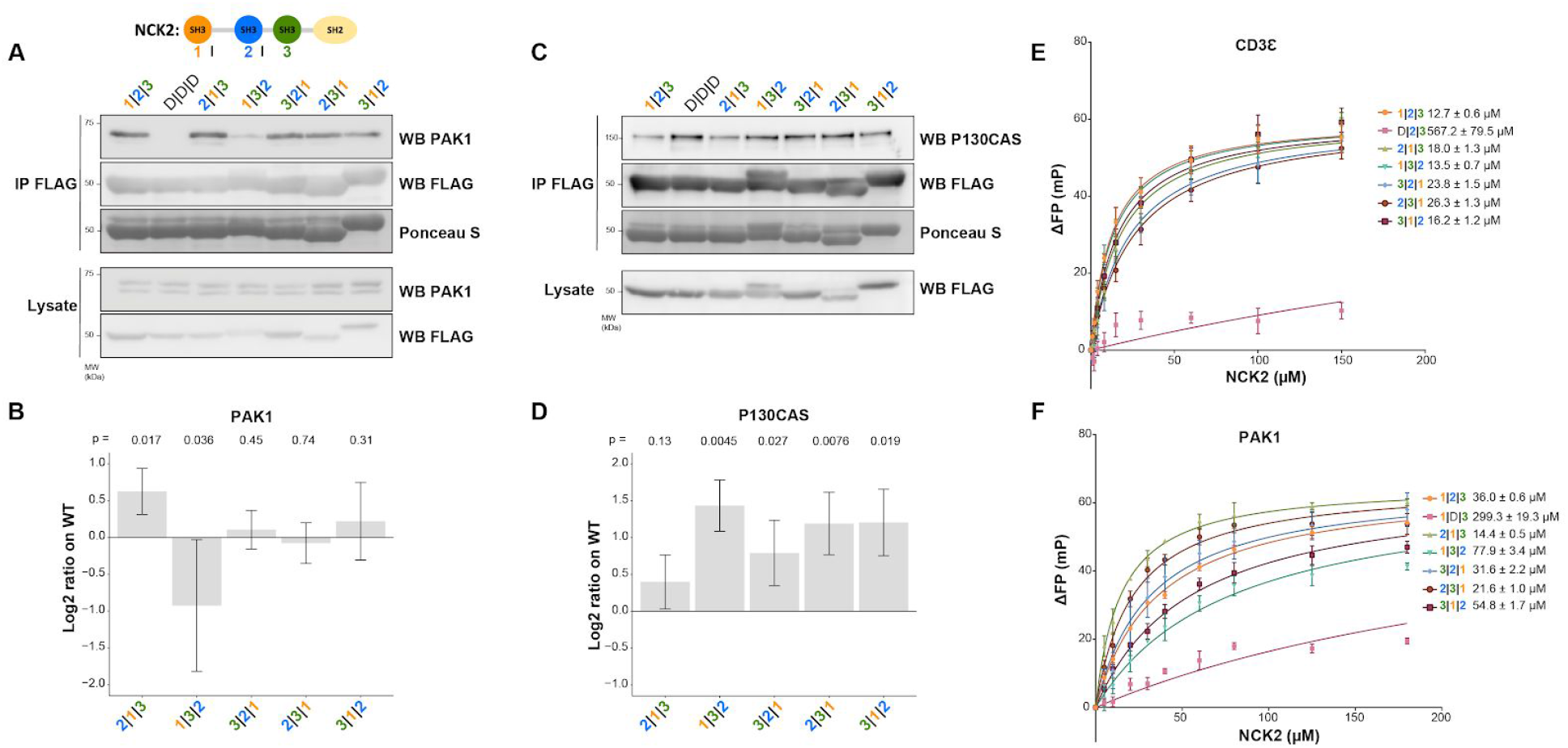
NCK2 SH3 shuffling affects its interactions with PAK1 and P130CAS/BCAR1 in cells but only slightly alters the availability of the SH3 binding pockets *in vitro*. A) Western blot analysis of NCK2 SH3-shuffled proteins interaction with PAK1. Each bait is N-terminally fused to a 3XFLAG tag. Loading corresponds to Ponceau staining of the nitrocellulose membrane. Experiment was performed in four replicates in HEK293T cells. B) Western blot quantification of PAK1 co-immunoprecipitation with NCK2. Average log2 ratio relative to the WT protein is shown (NCK2mut/NCK2WT, WT NCK2 log2 ratio = 0). Ratios from replicates were compared to WT NCK2 via pairwise ANOVA statistical test (p-values: 2|1|3 = 0.017, 1|3|2 = 0.036, 3|2|1 = 0.45, 2|3|1 = 0.74 and 3|1|2 = 0.31). Error bars indicate the SD. C) Western blot analysis of NCK2 SH3-shuffled proteins association with SH2 target P130CAS/BCAR1. The experiment was performed in four replicates in HEK293T cells stimulated with the tyrosine phosphatase inhibitor pervanadate. D) Quantification of the Western blot signals of P130CAS co-immunoprecipitation with NCK2 as in B). The four ratios of the replicates were compared to WT NCK2 via pairwise ANOVA statistical test (p-values: 2|1|3 = 0.13, 1|3|2 = 0.0045, 3|2|1 = 0.027, 2|3|1 = 0.0076 and 3|1|2 = 0.019). Error bars indicate the SD. E-F) Fluorescence polarization *in vitro* curves for NCK2 binding to CD3Ɛ or PAK1. Each binding assay was executed in triplicate. Error bars indicate the SD of the average of the ΔFP for each point. Dissociation constants error values represent the SE of the value derived from the binding curve. See also Table S6.

**Fig. S7.**
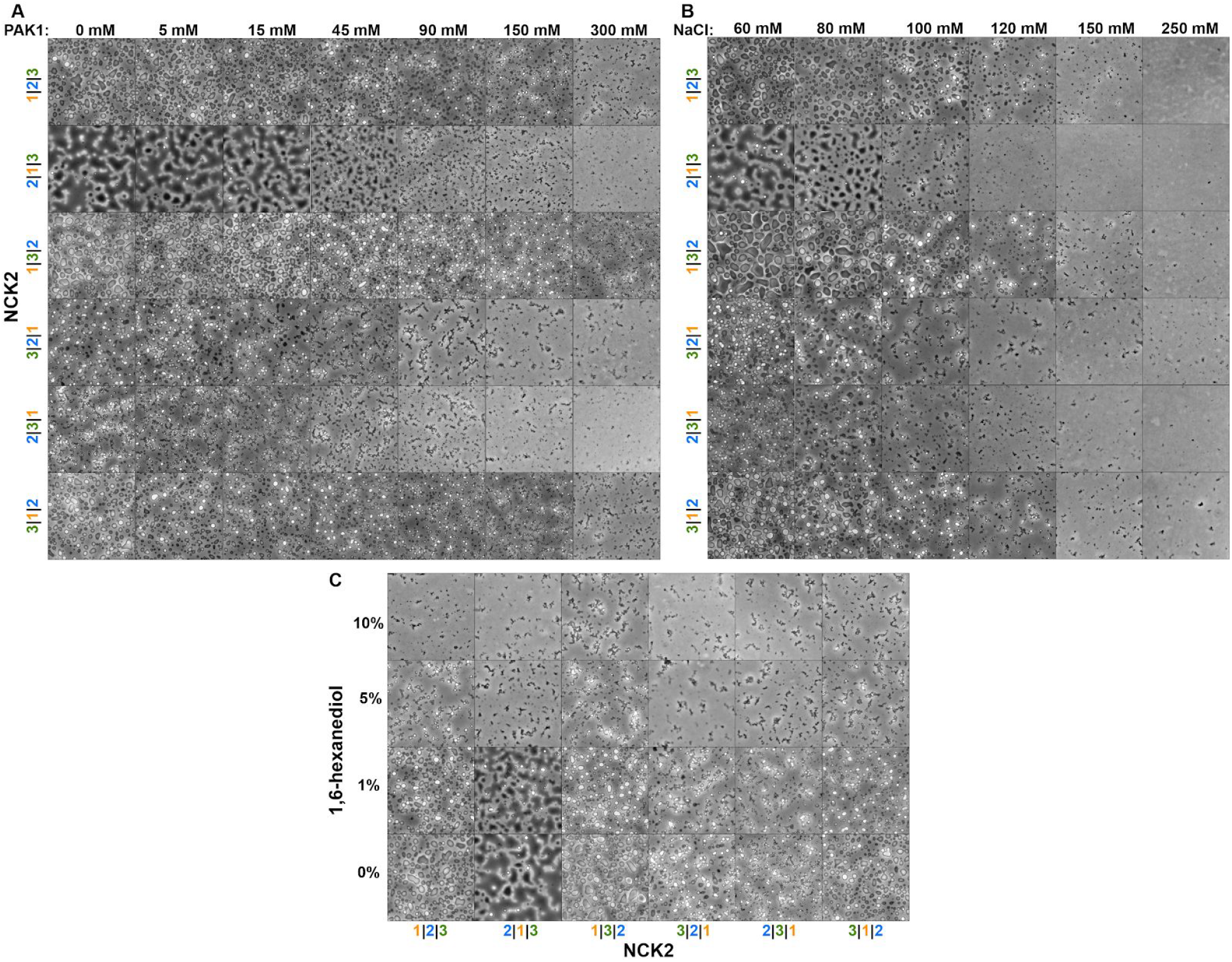
NCK2 SH3-shuffled proteins ability to promote phase transition is perturbed by PAK1, NaCl and 1,6-hexanediol. A-C) An increasing amount of PAK1 (A, PAK1 peptide), salt (B, NaCl) or 1,6-hexanediol (C) was added to the different NCK2 SH3-shuffled protein samples and incubated for 24 hours before observing phase transition using phase contrast microscopy. The experiment was performed in triplicates.

